# Genetic and Epigenetic Landscape of Self-Identified Hispanics/Latino in All Of Us

**DOI:** 10.64898/2026.10.30.763191

**Authors:** Adam C. English, Yilei Fu, Luis F. Paulin, Michal B. Izydorczyk, Sairam Behera, Qiuhui Li, Xinchang Zheng, Yiming Zhu, Justin Paschall, Grace E. Tietz, Jessica Honorato-Mauer, Luke B. Morina, Sean Griffith, Aishwarya Pisupati, Siyu Wang, Shannon Dugan-Perez, Helen Shen, George M. Weissenberger, Vanesa Vee, Jessica Gearhart, Michelle Kokosinski, Yi Han, Benjamin Myers, Alice M. Sanchez, Michelle Z. Mawhinney, Brian D. Craig, Beth A. Marosy, Heer H. Mehta, Kiran V Garimella, Ziad M. Khan, Eric Venner, Elizabeth G. Atkinson, Richard A. Gibbs, Evan E. Eichler, Michael E Talkowski, Kimberly F. Doheny, Donna Muzny, Michael C. Schatz, Ginger A. Metcalf, Winston Timp, All of Us Research Program Long-Read Working Group, Fritz J. Sedlazeck

**Affiliations:** Human Genome Sequencing Center, Baylor College of Medicine, Houston, TX, USA; Department of Computer Science, Johns Hopkins University, Baltimore, MD, USA; Johns Hopkins Genomics, Department of Genetic Medicine, Johns Hopkins University School of Medicine, Baltimore, MD, USA; Department of Molecular and Human Genetics, Baylor College of Medicine, TX, USA; Program in Medical and Population Genetics, Broad Institute of MIT and Harvard, Cambridge, MA, USA; Department of Genome Sciences, University of Washington School of Medicine, Seattle, WA, USA; Howard Hughes Medical Institute, University of Washington, Seattle, WA, USA; Department of Neurology, Harvard Medical School, Boston, MA, USA; Center for Genomic Medicine, Massachusetts General Hospital, Boston, MA, USA; Department of Biomedical Engineering, Johns Hopkins University, Baltimore, MD, USA; Department of Computer Science, Rice University, 6100 Main Street, Houston, TX, USA

**Author notes:** These authors contributed equally to this work (Adam C. English, Yilei Fu, Luis F. Paulin).

## Abstract

Human genetic studies still underrepresent many populations, limiting discovery and interpretation of medically relevant variation. Extensive admixture in self-identified Hispanic/Latino populations increases the need for representative reference data and ancestry-aware interpretation of medically relevant variation. We used long-read sequencing to analyze genetic variation, ancestry, and DNA methylation in 1,490 self-identified Hispanic/Latino participants from the All of Us Research Program. We identified 10 million novel variants and more than 100 candidate phenotype associations, with fine-mapping prioritizing structural variants over nearby single-nucleotide variants in more than half. We linked 3,329 tandem repeats to DNA methylation, resolved widespread allele-specific methylation, and identified ancestry-linked regulation across paralogous genes. These results reveal previously inaccessible genetic and regulatory variation and show the value of long-read sequencing for diverse populations.

## Introduction

Hispanic/Latino populations, comprising over 650 million people globally (∼8% of the world’s population), encompass heterogeneous and culturally diverse groups with origins across the Western hemisphere. Hispanic/Latino identity is a self-identified ethnic designation and is distinct from genetic ancestry, which here refers to patterns of inherited genomic variation reflecting shared descent and is inferred from genetic data relative to reference populations. Despite their profound genetic, ancestral, and cultural diversity, they are often erroneously treated as a single homogeneous group in biomedical research(*1–3*). This oversimplification obscures critical differences in disease risk, drug response, and genetic architecture among subgroups with admixture from Amerindigenous, African, and European components(*3*). In the United States, Hispanic/Latino individuals currently represent 19.5% of the population(*4*), and are one of the fastest-growing yet least explored demographics in research, despite bearing a disproportionate burden of several chronic diseases, including diabetes, obesity, and certain cancers(*5*, *6*). Failing to account for a population’s genomic complexity not only perpetuates health disparities(*7*) but results in missed opportunities to deepen our understanding of disease biology. Genetic variants unique to or more frequent in particular populations have led to the discovery of previously unknown disease loci and pathways. For example, studies of Latin American populations uncovered a risk haplotype in the gene *SLC16A11* that helps explain increased prevalence of type 2 diabetes (T2D) in Mexico(*8*). This haplotype is common in individuals of Amerindigenous ancestry but rare elsewhere and its discovery illuminated a new metabolic pathway influencing diabetes susceptibility, which offers therapeutic potential for T2D patients regardless of their allelic makeup. This and other studies(*9*, *10*) highlight the importance of prioritizing genetic diversity in biomedical research to capture population-specific variants, regulatory mechanisms, and disease susceptibility that are often missed when studies focus narrowly on individuals of a single ancestry.

The All of Us (AoU) program aims to catalog the biomedical histories and genetic diversity of its participants across the United States population, facilitating deeper understanding of genetic variants associated with diverse phenotypes and medical risk(*11*). As a national biobank, AoU has enrolled >849,000 individuals with the aim of broad representation across the U.S., including populations who have historically been excluded from large-scale studies(*12*). The AoU initiative expands opportunities to study genetic variants and patterns across ancestrally diverse participants, increasing the inclusiveness of genomic data available for biomedical research. Initial long-read studies in All of Us demonstrated the utility of this technology for resolving complex genomic variation and, more recently, for population-scale structural variant discovery and association analyses in participants who self-identified as Black or African American(*13*, *14*). As part of this effort (*14*), we generated long-read sequencing (LRS) for 1,490 genomes of individuals who self-identified as Hispanic or Latino to investigate variant types (e.g., structural variants, tandem repeat mutations) and other genomic modalities (e.g., methylation) that are particularly underexplored(*13*).

LRS technologies, such as those developed by Oxford Nanopore Technologies (ONT) and Pacific Biosciences (PacBio), have revolutionized the study of genomic variation by enabling the detection of complex structural variants (SV), tandem repeat regions (TR), and haplotype phasing with unprecedented accuracy and resolution(*15*). These capabilities are critical for understanding the full extent of population-wide genomic diversity, as they reveal variants that are systematically missed by short-read sequencing. This is especially important for admixed and diverse populations that harbour large novel alleles ill-represented in the human reference genome and largely absent from other catalogs of genetic variation. By uncovering this previously hidden variation and its potential link to disease, LRS can provide deeper insights into the genetic architecture of complex traits and inform the development of more precise clinical diagnostics and therapies, ultimately advancing personalized medicine(*16*, *17*). Furthermore, LRS directly enables epigenetic sequencing without any additional molecular treatment(*18*). This feature overcomes limitations of common array-based or short-read sequencing approaches by allowing simultaneous profiling of all CpG sites’ methylation alongside genomic variants(*18*), including within repetitive elements, offering the unique opportunity to directly map genome-wide methylation quantitative trait loci (mQTLs). By jointly measuring genetic variation and DNA methylation in the same individuals, long-read sequencing enables systematic investigation of how complex alleles, including structural variants and tandem repeats, relate to epigenetic regulation across genetically diverse populations.

In this study, we analyze genetic and epigenetic variants across 1,490 genomes of individuals who self-identified as Hispanic or Latino representing diverse ancestry clusters across the U.S(*19*). Throughout the manuscript “Hispanic/Latino” designates a self-reported ethnicity category and is not used as a proxy for genetic ancestry(*12*, *20*). Variant accuracy (SNVs, Insertions/Deletions, tandem repeats, structural variants) is validated using 421 samples sequenced as technical replicates with both ONT and PacBio. We report population structures, key variants and their implications, and reveal sub-clusters within the U.S. self-identified Hispanic/Latino population to provide the first comprehensive catalog of variants in this group, regardless of type or size. This includes 10,345,818 novel SNV and InDels, 39,151 novel SV alleles in 81 ACMG-classified and other medically relevant genes. We further emphasize the importance of incorporating local ancestry, especially at medically relevant loci, to improve interpretation and risk assessment. Complementing the genetic findings, CpG methylation analysis reveals how all variant types, including tandem repeats, influence mQTLs, affecting gene regulation. We identify epigenetic regulation in several paralogous gene groups tied to population-specific variants, underscoring their role in genetic risk studies and providing insight into regulatory mechanisms. These findings establish a foundational resource for biomedical research and underscore the critical role of ancestry-aware analyses in understanding gene regulation, disease risk, and personalized medicine.

## Results

### Landscape of self-identified Hispanic/Latino population variants

We selected 1,490 self-identified Hispanic/Latino samples from the All of Us short-read catalog that maximize the representation of common variants in this population(*21*) (Methods). These samples were sequenced at 25x-45x, with a mean of 34-fold coverage of ONT R10.4 LRS (see Methods). The sequencing reads covered ∼94.4% of the GRCh38 human reference genome at 10x coverage or higher for all samples (**Supplementary Fig. S1a**). The N50 read length of the sequenced samples was 22.3 kbp which resulted in phase block N50 of 1.2 Mbp, sufficient to phase nearly all genes in the genome along with flanking regulatory sequences (**Supplementary Fig. S1b**). Additionally, 84 samples from the Human Pangenome Reference Consortium (HPRC)(*22*) were included as a comparison set.

We identified a total of 88,344,833 SNVs/InDels, with 36.9M (41.7%) singletons (i.e. Allele Count (AC) of 1, see **Supplementary Fig. S1e,f**). We observed a genotype (GT) missingness rate of ∼6.8% for SNP/InDel (**Supplementary Fig. S1c**), reflecting a minority of positions where genotypes could not be confidently assigned across samples. Small variants showed a genome-wide transition/transversion ratio of 2.01 (**Supplementary Fig. S1g**), in line with expectations(*23*). Additionally, the average small variant heterozygous to homozygous alternate (het/hom) ratio was 1.58 genome-wide.

Structural variant calling (SV, ≥50 bp genomic alterations(*24*)) was performed with a new optimized population SV calling pipeline involving *Sniffles*(*25*) for discovery, *Truvari*(*26*) for merging, and genotyping with *Kanpig*(*27*). This pipeline produced a total of 740,466 high-confidence SVs over 1,490 samples. Individuals contained an average 21,491 SVs (∼8,560 deletions, ∼12,930 insertions; **Fig. 1a; Supplementary Fig. S2**). The final filtered SV genotypes had a GT missingness rate of 0.16%, with only 4,387 (0.59%) SVs having ≥5% GT missingness across samples (**Supplementary Fig. S1c**). The het/hom ratio per-sample ranged from 2.33 at the 5th percentile to 3.49 at the 95th percentile (**Fig. 1b**). SVs were classified by sequence and location as involving either transposable elements (TEs) or tandem repeats (TRs), revealing that 8.2% of SVs contained TEs while 82.8% of SV occurred within TRs (**Fig. 1a**). Among TEs, SINE-type SVs (41,993 total; 39,237 outside TRs) were 4.2 times more common than LINE-type SVs (9,919 total; 9,087 outside TRs).

**Fig. 1.**
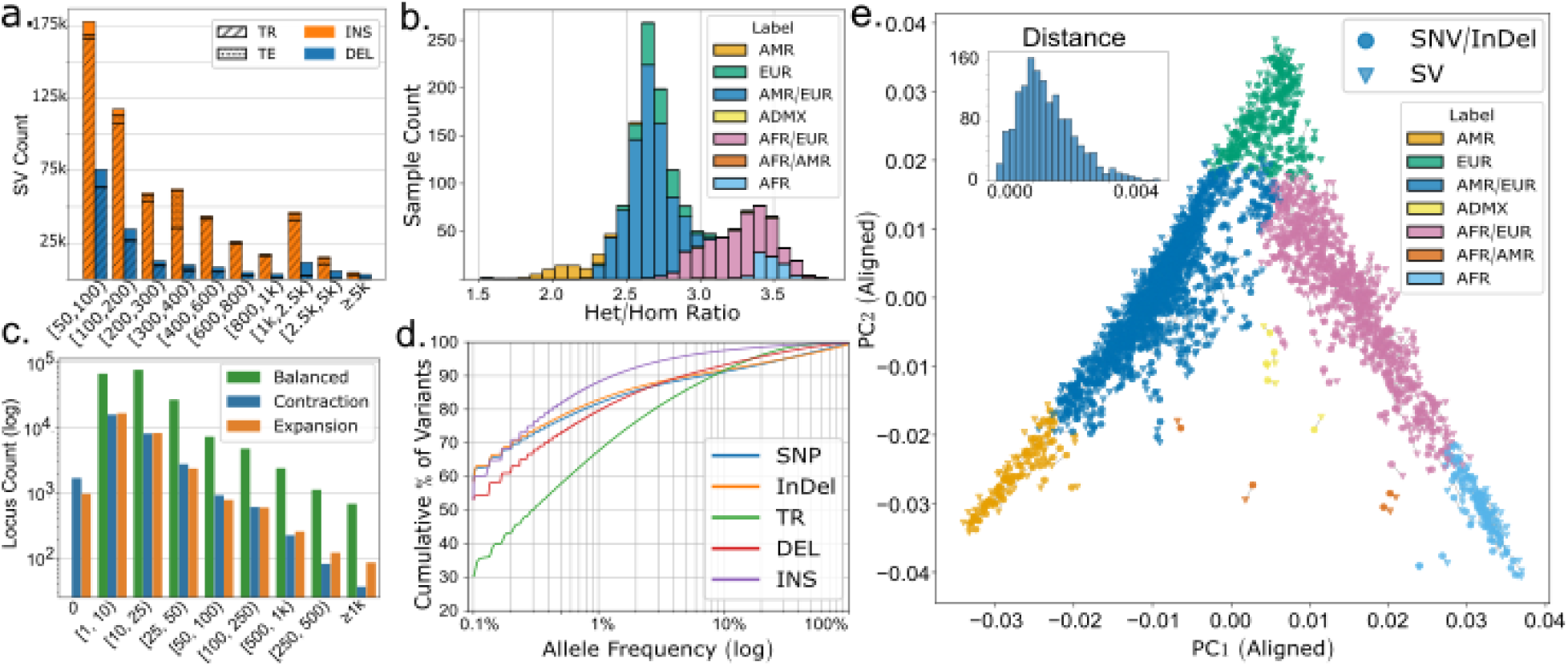
Variant Calling overview: **a.** SV type and size distribution with annotations of SVs occurring within tandem repeat (TR) or containing transposable elements (TE). **b.** SV Het/Hom ratio across Continental Ancestry Proportion (CAP) clusters. **c.** Tandem repeat calling allele length range across 302,175 regions (≥1% AF). **d.** Allele frequency distribution across all variant classes. **e.** Principal component analysis (PC1 and PC2) of SNVs/InDels and SVs, projected onto each other using Procrustes analysis. Samples are colored by CAP clusters. Interior histogram contains the Euclidean distance distribution in PCA space of samples’ SNP/InDel *and SV PCs*.

As SVs are predominantly within TR, and InDel calling within TRs is particularly difficult, we also performed TR specific variant calling for a more precise characterization in these regions (see Methods). This evaluated >300K TR loci, which had >33 million TR alleles across the population. TR alleles were also counted by their length resulting in 6,509,007 unique-by-length TR alleles (**Fig. 1c**). On average, each locus produced 19.3 unique-by-length TR alleles with 18,279 highly-variable loci having ≥50 alleles. A total of 1,999,273 alleles over 302,175 loci have ≥1% allele frequency by length. The length range of common TR alleles was defined as the difference between the minimum and maximum allele deltas (allele length minus reference length) at each locus. Most variability in TR allele length per-locus is balanced, meaning the locus has both expansion and contraction alleles. About half of the loci (162,065 loci, 53.6%) have a TR allele length range of ≥10bp, which is expected given the highly polymorphic nature of TRs(*22*). These loci with common (≥1%AF) and highly variable allele lengths (≥10bp) contain 1,537,382 alleles in total.

After variant processing, the overall allele frequencies (AF) of all variant types were found to exhibit the expected distributions of common and rare events (**Fig. S1d**). SVs across the cohort, for example, were dominated by singleton (18.2% of SVs) and polymorphic SVs (AF ≤ 50%; 80.9% of SVs) with only 0.76% of SVs being major alleles (50% < AF ≤ 100%) and 0.04% shared across all samples.

Of the 1,490 ONT samples sequenced, 421 were also sequenced as technical replicates using PacBio HiFi (mean coverage=34.8x; mean read length=17.1kb). These replicates were used to generate whole-genome, haplotype-resolved assemblies, from which variants were derived to serve as a baseline for benchmarking each class of discovered variants (see Methods). SNVs had an average F1 score (harmonic mean of precision and recall) of 0.975, InDels 0.634, SVs 0.917, and TRs 0.879 (**Supplementary Fig. S1d, Table S1, Supplementary Section 2)**. This benchmarking determined a per-sample precision/recall of 0.977 / 0.974 for SNVs and 0.951 / 0.886 for SVs. Furthermore, the average genotype concordance of SVs was 89.7%, and TRs was 93.5%. We also observed that the lower performance of InDels was explained by suppressed recall of 1bp to 4bp events, a known issue for ONT sequencing, which recovered to expectations for InDels 5bp-50bp (F1 0.735)(*28*) (**Supplementary Fig. S3a**). The performance of TR calls <50bp with F1 0.855 was higher than InDels within TR regions at F1 0.649. Similarly, the F1 of variants ≥50bp produced by the TR caller was 0.937 compared to 0.920 for SVs within TR regions. Overall this highlights the accurate variant calling and filtering across the 421 technical replicates.

The extensive and comprehensive nature of the SV call set allowed for an investigation into potential patterns among the remaining false positive (FP) SVs across different samples (**Supplementary Fig. S3b**). We hypothesized that FPs might arise at either high frequency (e.g., due to systematic alignment artifacts) or at low frequency (e.g., from stochastic noise). Supporting the latter, we observed a significant skew towards rare events (1-10 individuals; chi-squared test, p < 0.01), indicating that FPs are more likely to occur at low AF.

Variants were next evaluated in the context of population structure using principal component analysis (PCA) to assess whether they recapitulate expected population structures(*29*). This was performed independently for SNP/InDel and SV (**Fig. 1e** and **Supplementary Fig. S4a-c**), after batch correction of the SV PCAs attributable to the use of mixed basecaller versions (**Supplementary Fig. S4d**). We observed that each variant class’ PCA was largely consistent across PCs 1-3 revealing the same population structure. A Procrustes analysis measured the disparity between SNP/InDel PCs and SV PCs 1-3 at 0.061 (**Fig. 1e**). Furthermore, PCA was repeated on small variants beside a reference panel of 4,091 samples from a combination of the Human Genome Diversity Project (HGDP)(*30*) and the 1000 Genomes Project (1kGP)(*23*) (**Supplementary Fig. S4e**) which enabled us to profile the global representation of these individuals’ genetic diversity. This demonstrates that both SNVs and SVs consistently capture the diverse population structure within this cohort.

### Genetic ancestry within the self-identified Hispanic/Latino cohort

The above principal component analyses (**Fig. 1e**) revealed a continuum of genetic variation across this self-identified Hispanic/Latino cohort, consistent with complex admixture(*12*). To examine the cohort’s genomic composition, we next broadly grouped individuals by global continental ancestry proportions (CAP) and characterized individual haplotypes via local ancestry inference (LAI).

To measure CAP, ADMIXTURE(*31*) with K=3 was run using 7.6 million common (≥5% AF) genome-wide SNP/InDels and a reference panel of HGDP and 1kGP samples having known super-population labels of African (N=790), American (N=563), and European (N=749) (**Fig. 2a**; see Methods). Although an extensive reference panel was available, we opted for unsupervised ADMIXTURE analysis as we observed that its supervised mode with lower values of K was insufficiently sensitive to samples with high admixture (**Supplementary Fig. S5, S6**).

**Fig. 2:**
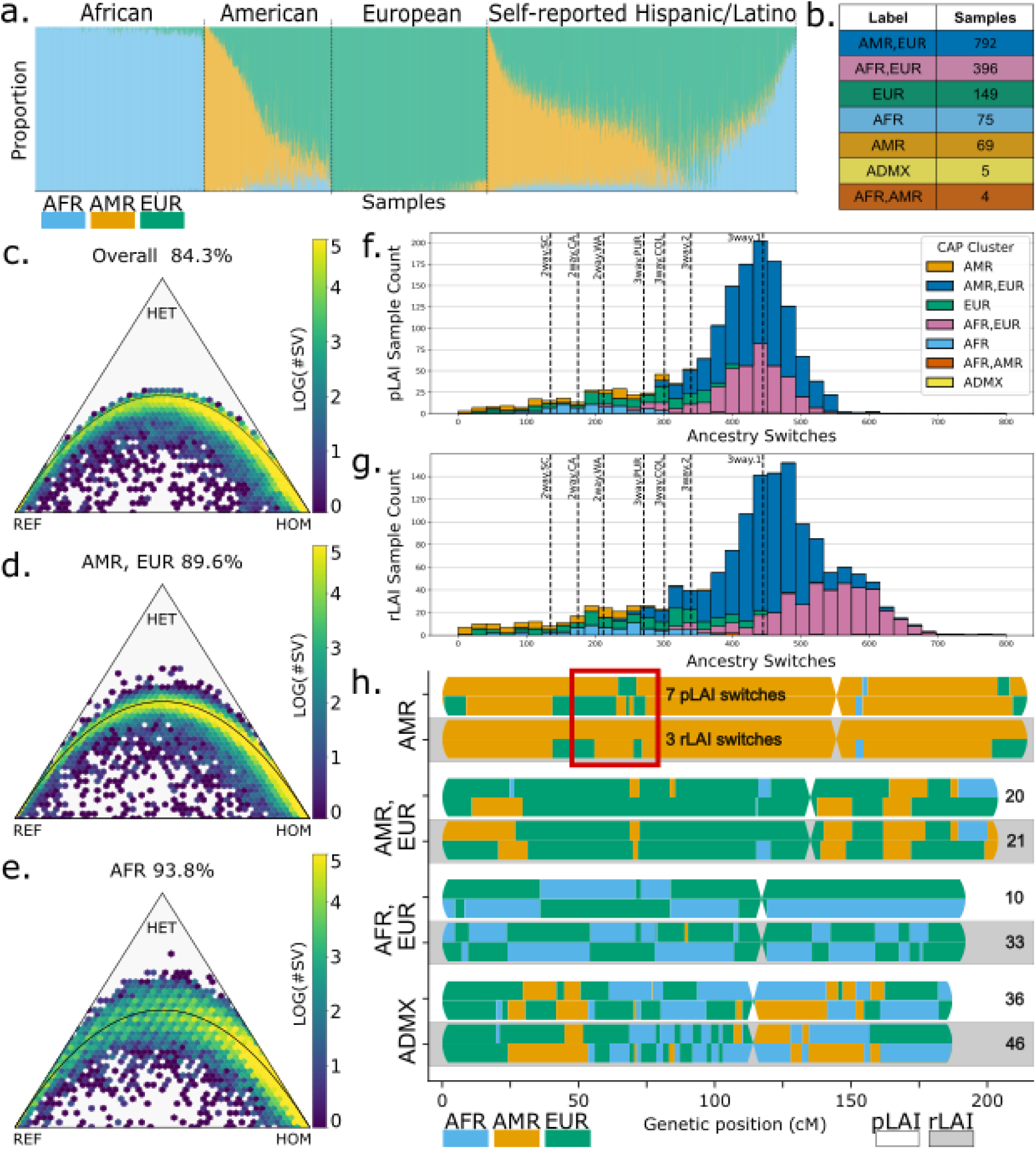
Population structure of self-identified Hispanic/Latino individuals of All of Us: **a.** Admixture plot showing continental ancestry proportion from ADMIXTURE with K=3 on 2,102 HGDP+1kGP short-read reference samples and 1,490 AoU samples. **b.** CAP (Continental Ancestry Proportion) cluster sample counts. **c.** Overall Hardy-Weinberg equilibrium (HWE) pass rate for SVs across all samples. Color represents the log SV count **d,e.** HWE rates for SVs within AMR/EUR and AFR CAP clusters **f.** Local ancestry switch count for population phasing **g.** Local ancestry switch count for long-read based phasing. For f and g, one outlier with >1,000 switches is not shown. **h.** Karyogram of four individuals’ pLAI and rLAI switches across haplotypes for chromosomes 4-7 with example switch counts (red box) and total switch counts (right).

Samples were assigned to a CAP cluster of genomically similar individuals based on the continental ancestries that accounted for at least 75% of the sample’s total CAP (**Supplement Fig. 7a).** For example, a 2-way admixed sample with 90% Amerindigenous (AMR), 10% European (EUR) received the AMR label corresponding to the strongest ancestral component whereas a 3-way admixed sample with e.g., 60% EUR, 30% African (AFR), 10% AMR received the dual label AFR/EUR. This resulted in 40 samples with lower levels of admixture and a single ancestry explaining ≥95% of their CAP, while 253 (16.9%) exhibited 2-way admixture, and 1,192 (80%) 3-way admixture (**Fig. 2b**). The 3-way admixed samples’ primary ancestry component explained an average of 57.5% of their genome and the second component 35.4%. Additionally, four samples had no two ancestries explaining 75% of their CAP and were labeled as ADMX. These assignments were validated using ancestry informative markers (AIMs) from two studies designed to assess CAP in highly admixed samples across the Americas(*32*, *33*). Compared to the genome-wide variants, this set of 540 AIMs produced highly correlating CAP (Pearson r > 0.98 for AFR, AMR, and EUR components; N = 1,490) and highly concordant clusters with 95.5% of samples consistently assigned (**Supplementary Fig. S7b,c**).

The importance of CAP clustering was apparent across several measures of population consistency. First, we assessed the Hardy-Weinberg equilibrium (HWE) of SVs, where a higher passing rate indicates lower population structure discordance. HWE testing across all samples collectively revealed a pass rate of 84.3% (α=0.05; **Fig. 2c**). However, this increased when stratifying by CAP cluster, ranging from 89.6% in AMR/EUR to 93.8% in the AFR cluster (**Fig. 2d,e**; **Supplementary Fig. S8**). CAP clusters also exhibited a tighter distribution of het/hom ratios compared to the overall distribution (**Fig. 1d**). Using F_ST_ as a measure of population differentiation, we compared the proportion of high-F_ST_ variants (>0.15) for both SNVs and SVs for all CAP cluster pairs excluding ADMX **Supplementary Fig. S7d**). We observed a higher proportion of high-F_ST_ variants between AFR and AMR clusters (22% for SNVs, 10% for SVs), with EUR showing intermediate proportions (11.02% for SNVs, 4.37% for SVs). Together, these measures demonstrated that CAP clustering grouped genetically similar individuals and captured meaningful population structure.

Due to the highly admixed nature of Hispanic/Latino individuals, sample-level, global ancestry assignments will not fully capture the complexity of ancestral haplotype structure, which can be especially important around medically relevant genes(*34*). To address this, we employed local ancestry inference (LAI) to assign ancestry at the haplotype level using two variant phasing techniques (see Methods). The first was population-based phasing (pLAI) from *Shapeit*(*35*) and a reference panel of 1kGP (see Methods). The second used read-based, individual phasing (rLAI). Genome-wide, both phasing strategies produced ancestry proportions that correlated highly to the global CAP (PearsonR >0.97; p<0.01).

As local ancestry allows for the identification and visualization of haplotype ancestry switching, we examined the distribution of local ancestry tracts in our empirical data to that of several simulated datasets. More continuous local ancestry blocks (i.e. longer tract N50 and fewer switches between ancestries within a haplotype) are generally indicative of higher quality results after accounting for switches that are expected due to admixture(*36*). Both phasing approaches produced ancestry tract N50s averaging ∼27 Mbp. The average number of ancestry switches per-sample in the AoU cohort was 390 to 428 for pLAI and rLAI, respectively (**Fig. 2f, 2g**). To contextualize these switch counts, we simulated ancestry breakpoints with varying ancestral haplotype proportions and generations of admixture for three levels of 2-way admixture and four levels of 3-way admixture(*37*) (**Supplementary Section 5**). Each CAP cluster’s level of admixture broadly corresponded to the number of switches expected. Specifically, predominantly 2-way admixed CAP clusters produced an average of 220 pLAI and 230 rLAI switches, consistent with 2-way simulations ranging from 135 to 212 switches. The 3-way CAP clusters (e.g., AMR/EUR) produced an average of 431 pLAI and 477 rLAI switches, and 3-way simulations between 271 and 445 switches. Both phasing techniques produced generally consistent ancestry switch counts for samples from most CAP clusters. However, the degree to which the LAI results agreed was negatively correlated to samples’ admixture entropy(*38*, *39*) (PearsonR -0.33; p <0.01; **Supplementary Fig. S9a-c**). For example, AMR samples had an average admixture entropy of 0.37 and an average of 25 more switches in pLAI than rLAI whereas AFR/EUR averaged admixture entropy of 0.91 and 109 fewer pLAI switches than rLAI. **Fig. 2h** illustrates this trend over chromosomes 4-7 for four representative samples across CAP clusters.

To corroborate these results, we also performed LAI using the haplotype resolved PacBio assemblies (aLAI; **Supplementary Fig. S9d,e**). These 421 samples were a representative subset of all CAP clusters (chi-squared = 0.020, p = 0.999) and allowed us to independently assess LAI. Across all CAP clusters, aLAI switch counts were more concordant to rLAI than pLAI with mean differences of 3 and 51, respectively. These results demonstrate particularly promising outcomes from the rLAI approach. While pLAI better fit the simulated ancestry switch counts, rLAI and aLAI, which used orthogonal sequencing reads from ONT and PacBio, respectively, agreed with one another. Both rLAI and aLAI reported a higher number of ancestry switches than pLAI for samples from CAP clusters with strong AFR components, suggesting that simulations and pLAI calculations may under-estimate ancestry switches for these genomes.

### Identification of transient variants across CAP clusters

Given the high quality of our SV call set together with the high resolution of LAI we are able to accurately assess whether specific mutations deviate from the general CAP cluster structure, potentially representing transient variants(*40*). Such transient variants could potentially highlight regions under selection or could reflect independently gained *de novo* mutations. To identify transient variants, we searched for SVs present in a single individual from one CAP cluster that are widely common in other CAP clusters (see **Methods**). For this, we implemented a new method to visualize and study transient variants. Briefly, for each SV each continental ancestry component (AFR, AMR, EUR, within CAP clusters or LAI) is summarized over all carrier individuals and projected into 2-D space. **Supplementary Fig. S10a** shows the ancestry composition of a given SV computed from the CAP cluster (purple circle) and pLAI (red circle), compared to the averaged ancestry composition over all individuals in the cohort based on CAP clusters (AFR: 17.5%, AMR: 31% and EUR: 51.5%) denoted as Null (black dot). This shows that the AMR and EUR composition decreases while the AFR increases for both CAP clusters (AFR: 33.3%) and pLAI (72.2%). Using this technique, we detected 31 transient SVs in 30 individuals (one individual from the control panel, not used in further analysis) with one individual from the cohort carrying two transient SVs (**Table S2**). Over half the transient SVs were within TR regions and four were TE-mediated, arising as a consequence of transposon activity. The 30 transient SVs overlapped with five medically relevant genes, one of which is related to male infertility (*RNF212*), one to the immune system (*FLT4*), and two to neural system functions (*ELP1, COL17A1*).

To further delineate if these variants occur because of recent admixture, we examined their distribution across local ancestry components. Out of the 30 transient SVs, 23 can be fully explained with pLAI, highlighting the importance of local vs. genome wide ancestry when studying alleles (**Supplementary Fig. S10b**). Nevertheless, three out of the 30 transient SVs can be only partially explained by pLAI while four SV cannot be explained (three INS, one DEL, **Table S3**). LD analysis shows ¾ SVs have no related SNPs (nor SVs) 1Mb up and downstream, while 1 SV has only 2 SNPs with r2 > 0.5. Notably, this SVs is linked to eight SVs (r2 >0.8) and an additional SV with r2 >0.5. Further, these four transient SVs are within TR regions and two genes might be affected: *PLEKHG4B* and *COL17A1*. The four SVs, which also are TR, show two major continental ancestries when analyzing the local ancestry of all carrier individuals (i.e. AMR and EUR) and are present in the HPRC controls from EAS and SAS ancestry.

Recent reports have shown that TE-mediated SVs (i.e., recombinations of two repeats e.g., ALU-ALU) can produce low frequency mutations where the individual SVs might be independently gained across individuals(*25*). To explore these, we investigated TE-mediated SV present across two or more CAP clusters focusing on rare SV in the self-identified Hispanic/Latino population (AF<1%). Among reported SVs, we found 84,347 TE-associated SV (e.g., ALU insertions), 55,199 INS containing variants and 25,899 TE-mediated DEL (3.50% of total SVs), from which 33.79% correspond to Alu-Alu recombinant and 18.17% to recombinant LINE/L1-LINE/L1 events (**Table S4, Supplement Section 1**). The majority of TE-mediated (19,480 SV) SV are rare (AF<1%) in the population. We performed LAI based ancestry composition summarization of rare TE-mediated SVs that had at least five individuals (4,835 SVs, see **Methods** for details). These SVs were then sorted by their distance to the population average with the aim of selecting SVs whose ancestry composition does not follow ancestry patterns the 4,835 rare TE-mediated SVs across CAP clusters (**Supplementary Fig. S10c)** and pLAI (**Supplementary Fig. S10d**). The CAP cluster ancestry summarization shows that most of the SVs have a strong AFR and EUR component (yellow-orange colored dots between the AFR and EUR nodes), with a second cluster of SVs surrounding to the population average (shown as a yellow shadow) and a third in the AMR/EUR (single yellow dot between the AMR and EUR nodes). In purple we highlighted the 100 SVs closest to the population average (black dot) based on LAI. Next, LAI ancestry summarization shows a clearer separation of the continental ancestries, with most of the SVs having predominantly AFR ancestry (yellow-orange colored dots close to the AFR node) with a second cluster of SVs close to the EUR node (yellow shadow). In red we highlight the same 100 SVs closest to the population average (black dot), which are tightly concentrated around the Null. Manual curation of these 100 SVs was performed to remove possible artifacts. **Table S5** shows the ancestry components of ten manually curated transient SVs. Two of these manually curated SVs have an ACMG classification of variants of unknown significance (VUS), while the other eight are benign. Five of the curated SVs overlap with introns of protein coding genes (*TRAPPC9, KCNIP3, ACAP2, KCNK10, SEPTIN7*) from which *TRAPPC9* is medically relevant. Furthermore, *TRAPPC9* and *SEPTIN7* have been associated with nervous system diseases(*41*) and *KCNIP3* has been associated with multiple diseases ranging from cardiac to neurological(*42*).

Overall we observe at least 30 transient variants across the self-identified Hispanic/Latino CAP clusters including SV and TR that are not solely explainable by pLAI structure. These variants are important for future studies as they disturb local and genome wide ancestry, which makes their interpretation harder despite impacting medically relevant genes.

### Identification of novel and clinically relevant variants

From these 1,490 LRS samples, we identified novel alleles and evaluated variants with existing clinical annotations within this AoU cohort. Of particular interest were variants classified as pathogenic or likely pathogenic, including heterozygous variants associated with recessive conditions that may be relevant to carrier status. To start, we evaluated the proportions of novel and known variants relative to other major catalogs of human variation, including dbSNP, gnomAD, and 1KGP (**Fig. 3a**; also see **Supplementary Fig. S11, S12** for region-wise distribution of functional impact). We identified 87.98 million small variants (SNPs and indels) on the autosomes. Of these, 10 million (11.9%) were not present in dbSNP157 and were deemed as novel. Furthermore, we identified 1,408,620 novel variants on the sex chromosomes. Nonsynonymous variants (N=509,538; 0.6%) were largely (90.42%) rare with allele frequency below 1% in gnomAD(*43*) and 1kGP(*23*) and approximately 90% of those were singletons. A total of 55,690 novel variants (0.06% of all variants) were located in exonic regions for all autosomes and 871 for sex chromosomes, representing 0.57% of all novel variants. Of the novel variants found in exonic regions, 37,423 were identified as nonsynonymous variants in autosomal regions and 629 for sex chromosomes (**Table S6**).

**Fig. 3.**
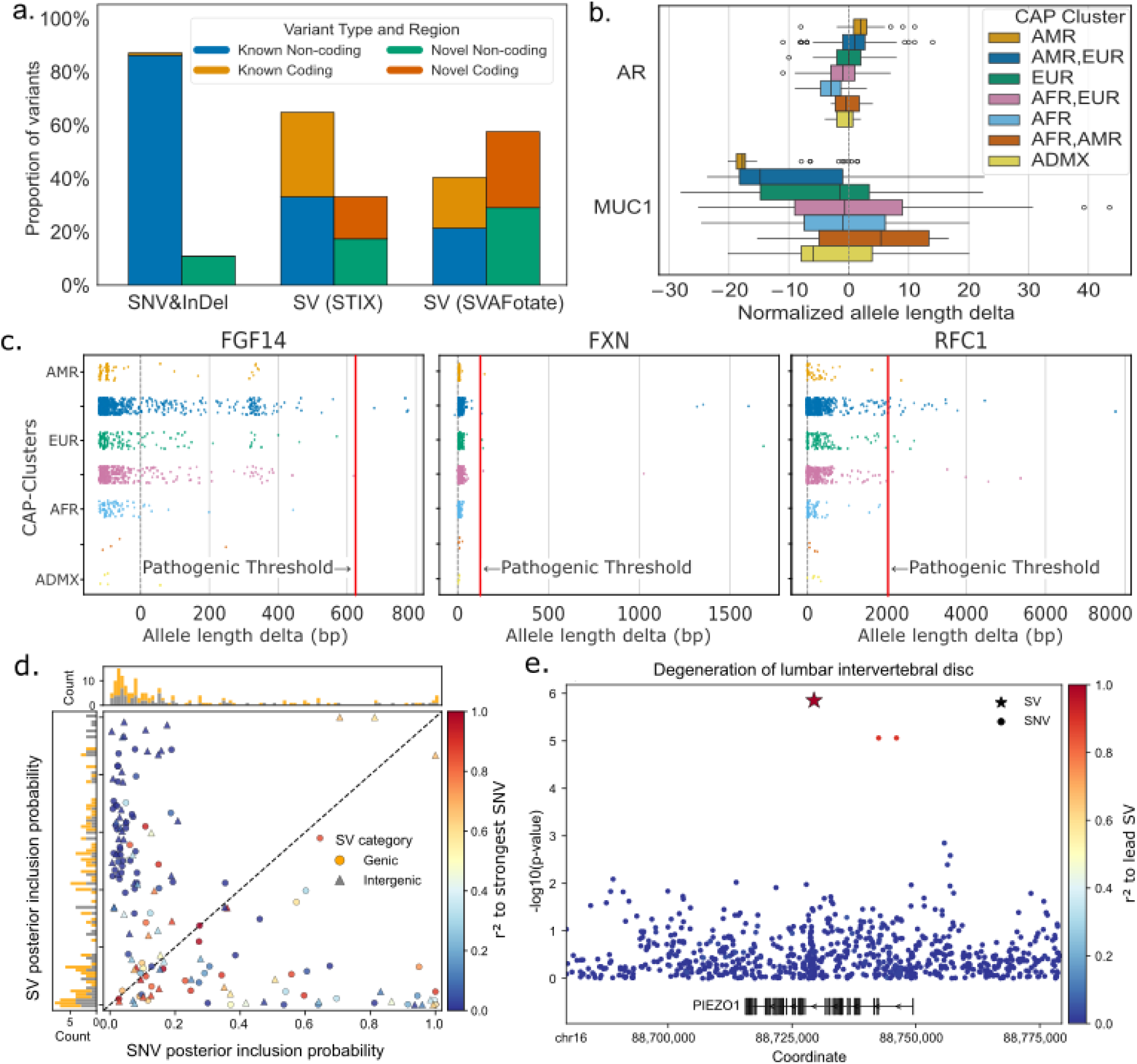
Novelty and impact of alleles across AoU. **a.** Rates of known and novel variants across small and structural variants. For SV we observed several genes hit by multiple thousand of SV, which skewed the measurement of coding impacting. **b.** CAP cluster specific examples of TR for AR and MUC1. Normalized allele length delta: TR length delta / motif length (boxplot lower quartile, 25th percentile; upper quartile, 75th percentile; center, median; extrema, 1.5 times the interquartile range). **c.** Detailed showcases of potential pathogenic TR alleles (indicated by red line). **d.** Comparison of posterior inclusion probabilities (PIPs) between each SV and the strongest nearby SNV (within ±100 kb) associated with the same phenotype after fine-mapping (central panel). Points are colored by LD (r^2^) between the SV and SNV. Circles indicate SVs within gene bodies, whereas triangles indicate intergenic SVs. The top and right panels show stacked histograms of PIP values for SNV-disease and SV-disease associations, respectively. Genic SVs are shown in amber, and intergenic SVs in gray. **e.** Manhattan plots showing a 681 bp insertion linked to degeneration of lumbar intervertebral disc. The insertion is the lead variant, and surrounding variants are colored based on their LD (r^2^) with the SV.

When comparing the SVs to four short-read derived SV catalogs(*44*) 63.25% of the AoU SVs are novel. This rate is inflated due to known limitations of short-read SV discovery(*45*). Therefore, we also assigned allele frequencies from STIX(*45*) utilizing 1,108 long-read genomes from 1kGP. STIX detected 39,151 (33.91%) novel SVs, serving as a more reliable assessment of Hispanic/Latino-specific SVs (**Supplementary Fig. S13**).

To identify pathogenic SNPs/InDels, we combined ClinVar classifications with review status higher than one star, REVEL scores ≥0.75, and CADD scores ≥20 (see Methods)(*46*). This uncovered 329 high-confidence pathogenic and likely pathogenic labeled variants with three present in more than 40 individuals (**Supplementary Fig. S14**; **Supplementary Section 7**) and 23 rare SNP/InDels located in ACMG-designated medically actionable genes. Consistent with a population-screening setting, these variants were primarily found in individuals presumed to be carriers rather than affected by disease. For example, four heterozygous pathogenic variants were identified in *ATP7B*, a gene associated with the recessive disorder Wilson disease. Separately, four singleton variants in *GAA*, a gene linked to Pompe disease(*47*), as well as three singleton alleles in *LDLR*, associated with familial hypercholesterolemia(*48*) Incidental carrier findings are not typically reported clinically unless specifically ordered.

Some potential pathogenic labeled mutations occurred at a higher frequency. For example, a heterozygous variant on the *HFE* gene (hereditary hemochromatosis(*49*)) occurred in 43 individuals (∼1.7% of population). This is a relatively common allele in European populations(*50*) and indeed the vast majority (31 individuals) of our samples show a European pLAI in this region. Furthermore, we identified a heterozygous variant in *BTD* that occurred in 58 individuals and is potentially associated with Leigh syndrome and Biotinidase deficiency(*51*). Local ancestry of this region revealed that the majority of individuals have European pLAI, but the variant also occurred in six individuals with African pLAI and nine with Amerindigenous pLAI. Additionally, variants impacting *G6PD* on chromosome X were found in 59 females with a heterozygous allele and eight individuals (six males and two females) with homozygous/hemizygous mutations. These variants are associated with *G6PD* deficiency(*52*) which, according to a recent screening study(*53*), occurs in ∼2% of the population.

One important strength of long-reads is their resolution of paralogous genes(*15*), such as *SMN1/SMN2* and *CYP2D6/CYP2D7*, in which certain variations are known to have phenotypic impact. Overall, 170 pathogenic labeled variants were identified in paralogous genes. One paralogous gene of note is *ABCA4,* which is associated with the recessive Stargardt disease and other retinal dystrophies in Hispanic/Latino populations(*54*). This gene contained 21 heterozygous pathogenic variants, with one occurring in 12 individuals and the others in fewer than 7, but since the associated disease is recessive, these individuals represent carriers. Similarly, nine heterozygous pathogenic variants (AC ≤ 2) were found in the *PAH* gene, which is associated with phenylketonuria(*55*).

We identified 215 (160 novel) high-risk candidate SVs (173 likely pathogenic, 42 pathogenic; **Table S7**) from which three could have potential functional consequences (see **Methods**). These SVs impact cardiovascular (*CD36, NPR3*) and metabolic (*GBA1, LPL*) genes, each with AC < 20, which have been previously reported(*56*) (**Supplementary Section 7.2**). Notably, labeled pathogenic SVs occurred in 31 individuals in *GBA1,* which is associated with Gaucher and Parkinson’s disease(*57*). Further, 26 individuals have a pathogenic-labeled SV in *HBA1.* Complex SVs in hemoglobin genes (*HBA1/HBG2*) may underlie reported *HbA1c* differences in Hispanic/Latino groups(*58*). Furthermore, two potentially functional SVs (homozygous DEL) affected the *MUC1* and *PLIN4* genes in two and four participants, respectively, while the rest were heterozygous carriers (14 and 111 individuals respectively).

We further filtered for novel SVs, identified using STIX’s population frequency annotation, and cross-referenced them with the ACMG clinical classifications (see **Supplementary Section 7.3** for details). Of the 250,703 novel SVs (33.86%), 114,973 were classified as benign, 38,802 as VUS, 128 as likely pathogenic, and 32 as pathogenic. The likely pathogenic and pathogenic SVs impacted 105 genes, including the aforementioned *GBA1 and PLIN4,* which are involved in lipid metabolism(*59*); *MUC1* which associated kidney disease(*60*); haemoglobin subunits *HBA1* and *HBG2,* reported to have differences in Hispanic/Latino groups (*58*) and *RIPK1,* which is associated with the auto-inflammatory syndromes(*61*).

We examined whether any CAP cluster-specific variants exhibited potential pathogenicity. For this, we identified variants from all variant types that have a F_ST_ ≥ 0.15 and are therefore relatively fixed in the CAP clusters (**Supplementary Fig. S7d**). For SNP/InDel we found 1,303,907 (1.48%) novel variants with 7,097 in exonic regions and 307,382 with allele frequency <1% in gnomAD and 1kGP. Remarkably, 791,686 SNP/InDel including 12,050 exonic and 256 loss of function (LoF) mutations reside in the 5,030 clinically relevant gene regions and 11,847 (286 exonic, four LoF mutation) SNP/InDel reside in the ACMG medically relevant gene list with 1,806 in the *RYR2* gene and 944 in the *TRDN* gene; mutations in both the genes are associated with risk of sudden cardiac death(*62*). We found 12.76% of SVs have a high F_ST_ (≥0.15) across CAP clusters with five deletions being classified as likely pathogenic overlapping four genes: *CCDC40, COL6A2, MUC1.* Notably, these highlight several paralogous genes such as *MUC* and *GUSBP4*. Furthermore, 641 SVs were classified as VUS and 6,030 as benign. The VUS-classified SVs showed an enrichment in nervous system development genes (105/307).

Specialized tandem repeat (TR) calling software enabled targeted analysis over 336,417 TR regions, including 186 TR loci previously linked to phenotypic and pathogenic impacts (*63*). Interestingly, these 186 loci showed greater length polymorphism than the broader TR set, underscoring the importance of screening these regions in particular (KS test p-value < 0.001**; Supplementary Fig. S15**). For 61 of the TRs, STRchive(*64*) provided annotations of TR-associated diseases’ pathogenic allele length thresholds. Analysis of these 61 loci by CAP cluster and allele length revealed 30 TRs with a significant association (one-way ANOVA, p<0.05), including *MUC1* and *AR* (**Fig. 3b**). These shifts across CAP clusters suggest population stratification of TRs that is potentially driven by demographic history, natural selection, or local adaptation.

A total of 135 individuals possessed expansions exceeding the STRchive annotated pathogenic allele length, including 8 TRs in which large expansions were somewhat common (AF ≥0.05%; **Fig. 3c; Supplementary Fig. S16** and **Table S8)** For instance, 10 individuals harbored heterozygous *FXN* alleles with >66 GAA repeats, exceeding the pathogenic threshold for autosomal recessive Friedreich’s Ataxia(*64*) (**Supplementary Fig. S17**). Five *FXN* alleles exhibited extremely large expansions (**Fig. 3c)**, with tract lengths approaching ∼1,700 repeats, consistent with previously reported disease-causing alleles(*64*). Additionally, expansions in the *FMR1* gene above the intermediate threshold of ≥45 repeats were found in 18 individuals and 5 individuals had one allele that reached the pre-mutation threshold of >60 repeats. No individuals possessed *FMR1* alleles surpassing the pathogenic threshold of ≥201 repeats(*64*) (**Supplementary Fig. S18**). While these alleles do not indicate individual participant risk, they do indicate the potential of increased risk for their offspring. A potential novel *RFC1* AATTG motif was also discovered in the Hispanics/Latino population. For some pathogenic TRs, such as *FGF14* (**Fig. 3c**), we observed that only individuals in the AMR_EUR CAP cluster exceeded the pathogenic threshold.

The only pathogenic TR locus with large expansions on both alleles in single individuals was within the Duchenne Muscular Dystrophy (*DMD*) gene(*64*). This TR currently has sparse and conflicting evidence linking it to *DMD*, a severe neuromuscular disorder with a median survival age of 27 years(*65*). Three females had homozygous *DMD* expansions above the pathogenetic threshold. For two individuals, the allele lengths reported from ONT data were confirmed using PacBio technical replicates’ assemblies and directly from the raw reads using TRGT(*66*) analysis. The third had no replicate. Furthermore, the clinical records of the 3 individuals reported no symptoms associated with *DMD*. These observations challenge the reported pathogenic interpretation of this *DMD* repeat expansion and highlight the importance of population-scale long-read data for refining variant pathogenicity annotations.

While pathogenicity of many TRs can be predicted solely by allele length, others are also influenced by motif composition. In the AoU samples, substantial expansions involving nonpathogenic motifs were observed at several disease-associated TR loci including *MUC1, BEAN1*, *SAMD12*, *YEATS2*, and *STARD7*. Although their motifs did not correspond to known pathogenic configurations, the repeats’ lengths still suggest potential locus instability and highlights the need for sequence resolved TR mutation studies.

Together, these findings demonstrate the power of long-read population sequencing to uncover rare and novel variants across all classes, with direct relevance to clinical and reproductive health. They also underscore the need for haplotype resolved, ancestry-aware genomics to enable precision medicine for all populations.

### Phenome-wide analysis identifies SV–phenotype associations

To investigate the clinical impact of SVs, we performed a phenome-wide association study (PheWAS) in 1,095 ONT participants with available clinical phenotypes derived from electronic health record (EHR) data in AoU CDR v8 (**Methods**). We identified 160 candidate SV-disease associations spanning 57 clinical conditions at a prioritization threshold of p < 1 × 10^-5^ (**Table S9**). Lower-frequency SVs tended to show larger effect sizes across prioritized associations (**Supplementary Fig. S19a**). We then conducted fine-mapping to identify SVs likely driving the disease associations while accounting for linkage disequilibrium (LD) with nearby SNVs. SVs were included in the 80% causal set for 139 (86.9%) associations and had the highest posterior probability in 96 (60.0%) associations (**Fig. 3d**), consistent with reports in self-reported African Americans of SVs being identified as candidate causal variants(*14*). One notable signal involved a 681 bp insertion in *PIEZO1*, a mechanosensitive cation channel gene previously implicated in intervertebral disc degeneration (*67*). The insertion was associated with lumbar intervertebral disc degeneration (odds ratio [OR] = 8.15, p = 1.41 × 10^-6^) and was the leading fine-mapped variant (**Fig. 3e**, **Supplementary Fig. S19b; Table S9**).

### Methylation diversity in self-identified Hispanic/Latino populations

DNA methylation plays a critical role in epigenetic regulation, but its relevance to medical phenotypes and disease remains underexplored compared to genetic or transcriptional profiling(*18*). Methylation patterns are relatively conserved across adult individuals within the same tissue, especially when compared to differences between tissues, reflecting the role of epigenetics in gene regulation(*18*). Despite the tissue-specific nature of methylation patterns, blood-based methylation provides a practical approach for studying methylation effects at population scales(*18*).

Using Nanopore-based methylation profiling the genome-wide DNA methylation rates (27,660,298 CpGs) across the 1,490 blood-based AoU samples revealed a strongly bimodal distribution with ∼74.7% of CpGs to have high DNA methylation above 80% and 8.8% of CpGs with methylation below 15% (**Supplementary Fig. S20a**). As a first QC step, we constructed a PCA of DNA methylation based on methylation frequency at each CpG (see **Supplementary Section 9.1**). As expected, PCA uncovered minimal DNA methylation patterns with PC1 explaining only 1.59% of the variance and largely capturing technical batch effect on the methylation levels created by divergent basecaller versions (**Supplementary Fig. S21**). Additionally, methylation PCA components 4 and 6 recapitulated the population structure observed from SNP/InDels (Procrustes disparity 0.2088) and SVs (0.2103) (**Fig. 4a**, **Fig. 1e**). We also investigated the per-chromosome PCA, where the actual epigenetic difference among the population can be more dominant compared to other factors (**Supplementary Fig. S22**). Differentially methylated CpGs between individuals comprised 770,828 methylation-variable regions (≥200 bp span; ≥2 CpG methylation levels; see Methods) that covered 10.47% of autosomal chromosomes (**Supplementary Fig. S20b**). ENCODE cis-regulatory elements (cCREs) from all tissues were next intersected with these methylation-variable regions. In total, 130,386 (16.9%) methylation-variable regions intersected cCREs, including 77,651 overlapping multiple cCREs. Enhancer-associated cCREs exhibited the strongest methylation variability, distal enhancer-like signatures (dELS) showed the highest absolute overlap (212,059 regions, 27.56% of all dELS), and proximal enhancer-like signatures (pELS) displayed the greatest proportional variability (68,281 regions, 40.65% of all pELS) (**Table S9**). CTCF-bound cCREs also showed substantial variation (155,000 overlaps, 35.14% of CTCF sites), whereas promoter-like signatures (PLS) had fewer variable regions (11,823, 29.85% of PLS). These intersections with ENCODE annotations were consistent with each active regulatory contexts’ roles in gene regulation(*68*), highlighting cCREs as key epigenetic variability hotspots and supporting their role in dynamic gene regulation.

**Fig. 4.**
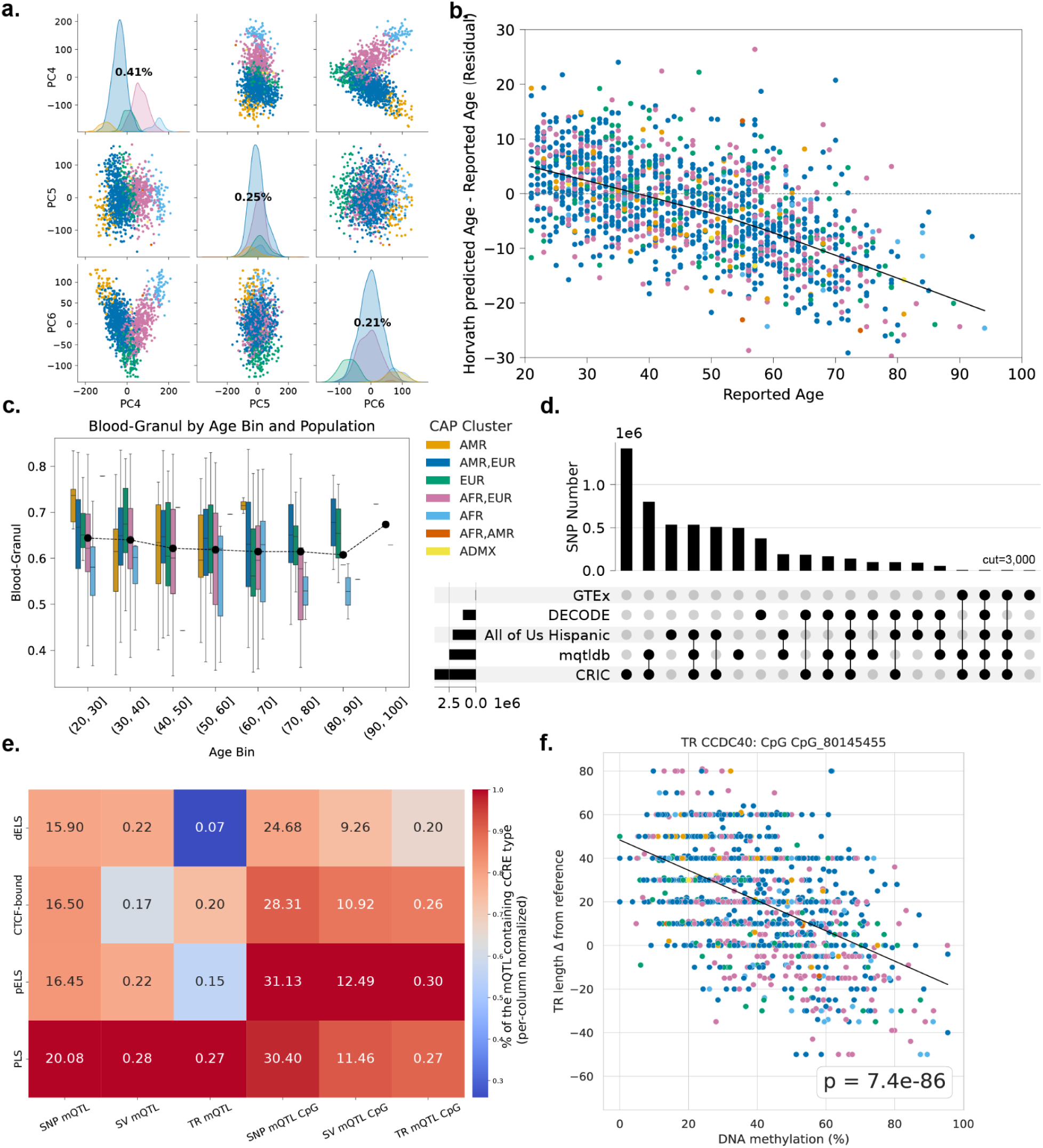
Methylation landscape. **a)** Pairplot of PC4 to PC6 where DNA methylation shows population stratifications. Diagonal: percent of explained variation for each PC. **b)** Residual plot between Horvath predicted age and reported age. **c)** Methylation-based granulocytes proportion estimation across CAP clusters. (boxplot lower quartile, 25th percentile; upper quartile, 75th percentile; center, median; extrema, 1.5 times the interquartile range). **d)** Comparison between AOU self-identified Hispanic/Latino mQTL SNPs to existing datasets. **e)** mQTLs and their associated CpGs’ overlap with ENCODE cCRE region **f)** An example of the tandem repeat on CCDC40 gene negatively correlating with a representative CpG in an upstream CpG island.

The comprehensiveness of this study’s available data with respect to methylation levels and self-reported age allowed us to compare participant’s self-reported age to samples’ epigenetic age prediction and observed a high correlation (Pearson r = 0.83; **Supplementary Fig. S23a**). Closer inspection revealed some age estimation deviations, which have also been observed in other studies(*69*). Specifically, individuals under 35 tended to have an overestimated predicted age whereas those over 40 tended to be underestimated (**Fig. 4b; Supplementary Fig. S23b**). Prior studies have suggested methylation age predictions are - in part - driven by cell type composition(*70*). Therefore, we performed cell type deconvolution to infer samples’ blood cell proportions using a reference methylation atlas(*71*) (outlier removed, see **Methods**). Granulocyte proportions decreased with age(*70*) (Pearson r -0.10; **Fig. 4c**) while T cells (r = 0.09) and monocytes/macrophages (r = 0.08) showed modest increases, consistent with known immunological aging dynamics (**Supplementary Fig. S23a**)(*71*). The increased proportion of granulocytes at older ages may reflect age-related myeloid bias associated with clonal hematopoiesis. Stratifying blood cell type deconvolution outcomes by CAP clusters revealed some associations. The AFR cluster contained moderate lower granulocyte cell type proportions compared to other CAP clusters (one-way ANOVA p<0.0001; **Fig. 4d**). Additionally, we observed CAP cluster dependent proportion shifts among T-cells and B-cells (**Supplementary Fig. S24**).

LRS enables joint measurement of methylation beside all variant types (SNPs, InDels, SVs, and for the first time TRs), thus allowing an integrated methylation quantitative trait locus in cis-windowed (cis-mQTL) analysis across all individuals. Overall, our Cis-mQTL analysis identified genetic regulation of DNA methylation from 2,456,683 regions that collectively covered 8.88% of genome-wide CpGs. A total of 1,366,276 (1.55%) SNPs and InDels, were significantly associated with 2,323,954 CpG sites (FDR corrected p-value < 0.05). We compared these mQTL hits and their associated CpGs with four independent datasets (see Methods). Comparing mQTL SNPs across all resources revealed that the majority were validated in these other works, but 532,314 (39%) of the AoU mQTL SNPs are novel relative to the other datasets, including 3,307 (0.62%) SNPs identified as novel by dbSNP157 (**Fig. 4d**, complete version at **Supplementary Fig. S25a**). We further investigated the uniquely reported mQTL SNPs from DECODE, the only other long-read study, and identified 27.75% of their mQTL SNPs were present in our study but not detected as mQTL SNPs in our cohort (**Supplementary Fig. S25c,d**).

Compared to the previous studies, our results produced a much higher number of mQTL SNP associated CpGs (2,323,954), with 79.56% being novel (**Supplementary Fig. S25b**). For GTEx, mQTLdb, and CRIC Atlas, this increase is largely due to methodological advantages provided by long-reads. While long-reads are capable of measuring all CpGs, these three prior studies leveraged DNA microarrays which only profile ∼2% of genome-wide CpGs(*18*). Additionally, our use of the newer ONT R10 chemistry and comprehensive variant discovery proved advantageous relative to the DECODE dataset.

Few studies have assessed the interplay of SV and methylation (SV-mQTL)(*72*). We identified 39,828 (4.66%) SVs with significant correlations to 830,270 CpGs. Furthermore, we jointly measure TR lengths and CpG methylation within the same native long-read sequencing assay to identify mQTL-TR relations. For TR-mQTLs, we associated CpG’s methylation level (M-value) against the TR length, which is different from traditional mQTL studies, which associate against variants’ presence/absence(*18*). For this, we selected 16,818 TRs from the top 5% of TRs with the largest length variability (Methods).

With the sets of mQTL variants and CpG sites derived across variant types, we next intersected each to ENCODE cCREs and measured each annotation types’ proportional burden as the percent intersecting an mQTL hit (**Fig. 4e, Table S10**). Promoter-like signature (PLS) cCREs were the most intersected by all variant classes with at least one SNP or InDel intersecting 20.08% of PLS, SVs intersecting 0.28%, and TRs intersecting 0.27%. Likewise, mQTL CpGs linked to all variant classes were also concentrated in PLS with SNP-CpGs 30.40%; SV-CpGs 11.46%; TR-CpGs 0.27%. Additionally, mQTL CpGs were most concentrated in pELS (SNP-CpGs 31.13%; SV-CpGs 12.49%; TR-CpGs 0.30%). These increased mQTL intersections in PLS and pELS relative to other cCREs support these elements being hotspots for DNA methylation changes.

In total, we identified 3,329 TRs that correlate with 17,714 CpGs. Out of these, 6 overlapped TRs previously reported(*63*) to have phenotypic impact: *KRTAP4-12, ZNF713, CCDC40, SSC5D, DRD4 and AC010336.3*. Among those six phenotypic TRs, *ZNF713* has previously been found to be correlated with DNA hypermethylation in a downstream CpG island, confirmed by our finding(*73*) (**Supplementary Fig. S26** and **Table S11**). Also, the *CCDC40* gene is negatively correlated with DNA methylation in an upstream CpG island (**Fig. 4f**).

TR expansions have been widely reported to alter CpG island methylation(*74*), and changes across these regulatory regions often reflect functional consequences of TR length variation. We identified 224 TRs with mQTLs linked to more than 2 CpGs in the same CpG island. A total of 129 of these TR-mQTLs were from TRs in proximity to but not overlapping the CpG island. We observed no general bias indicating stronger TR correlations to upstream or downstream CpG islands (50 and 46 TRs, respectively; **Supplementary Fig. S27**), nor significant difference in correlation coefficient magnitude was detected between the two orientations. These findings establish tandem-repeat length variation as a widespread correlate of local DNA methylation, extending the regulatory landscape of genetic variation beyond SNPs and structural variants.

Previous studies have highlighted the link of methylation with the expression of paralogous genes(*75*). Building on our discovery of novel mQTL associations, we examined whether paralogous genes contributed to the observed methylation patterns by analyzing SNP-CpG mQTL pairs located within genes from the same paralogous group (**Supplementary Fig. S28,** see **Methods** for details). A notable example was within the solute carrier family 22 (*SLC22*) paralogous group where variants on one copy of the gene impacted the methylation status of the other (**Fig. 5a**). Here, the SNPs chr5_132316836_A_C in *SLC22A4* (**Fig. 5b**) and chr5_132373185_T_C in *SLC22A5* (**Fig. 5c**) are strongly associated with methylation levels at chr5_CpG_132387043 in *SLC22A5* and chr5_CpG_132302683 in *SLC22A4* and their homozygous genotypes (CC and TT, respectively) are associated with CpG hypermethylation (p-value <0.0001 for both). Prior studies have shown that *SLC22A4* and *SLC22A5* are frequently co-expressed and involved in inflammatory disease risk loci, especially in the *IBD5* region(*76*), however these two specific SNPs have not been previously reported as mQTLs. Nevertheless, both variants are reported as eQTL variants in the GTEx eQTL database, where chr5_132316836_A_C (on *SLC22A4*) is a strong eQTL SNP for *SLC22A5* (p-value <0.0001 in whole blood), and chr5_132373185_T_C (on SLC22A5) is a strong sQTL (SplicingQTL) SNP for *SLC22A4* (p-value<0.0001 in whole blood). We further looked into the F_ST_ score of those two variants. The *SLC22A4* SNP differentiates the AMR and EUR CAP cluster (F_ST_ = 0.36), while the *SLC22A5* variant differentiates the AFR and EUR CAP cluster (F_ST_ = 0.37) (**Fig. 5c**). Interestingly, prior studies have actually shown the expression difference, which might occur due to methylation changes shown here in the European population in *SLC22A4* and *SLC22A5*. This highlights that CAP cluster specific variants could lead to activation or inactivation of different paralogous genes.

**Fig. 5.**
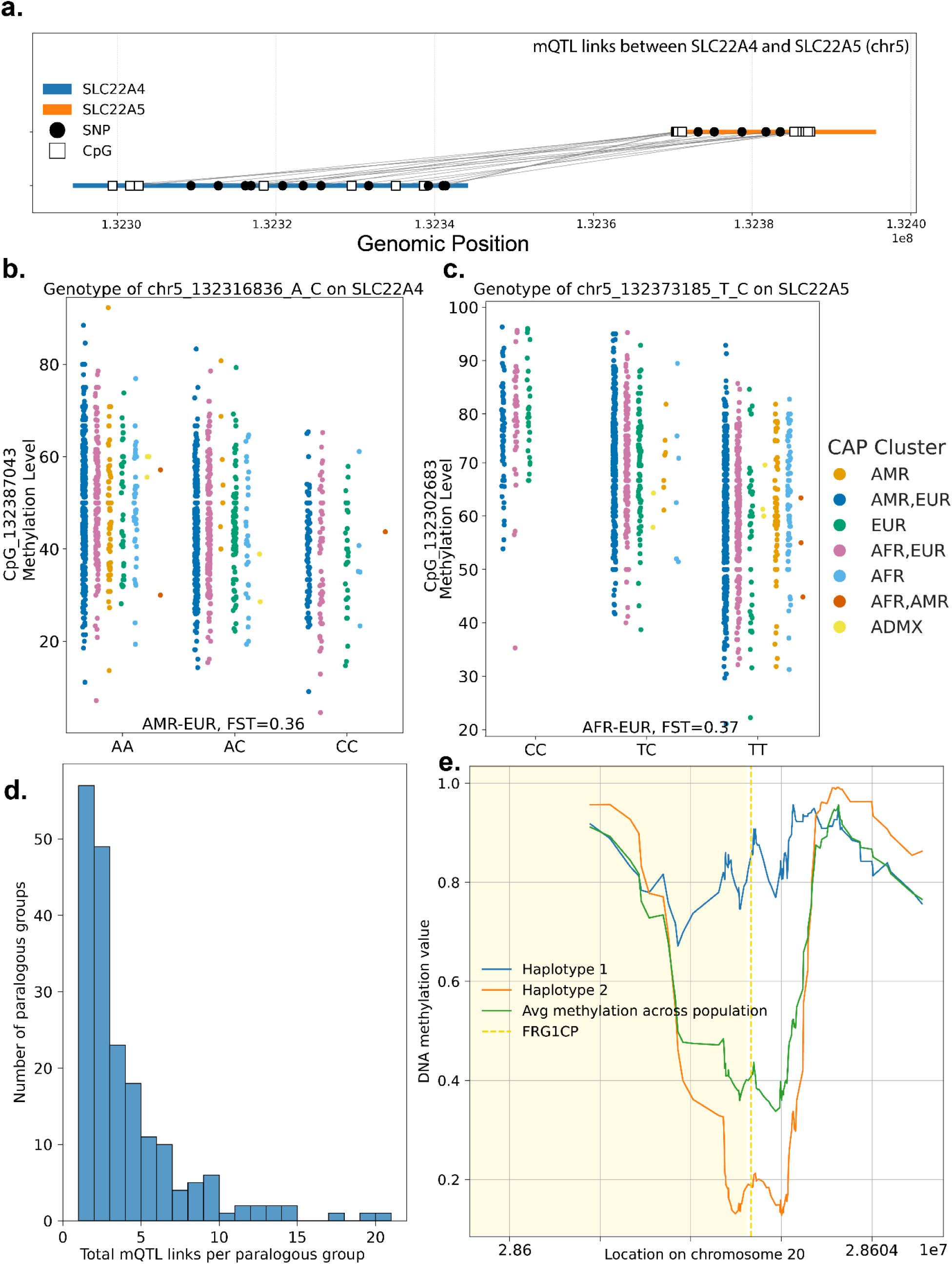
**a.** SNP–CpG associations from SLC22A5 to SLC22A4. **b.** Reciprocal mQTL co-regulation between the paralogous genes SLC22A4 and SLC22A5, where SNPs in one gene associate with CpG methylation in the other; **c.** Representative SNP–CpG associations from SLC22A4 to SLC22A5. **d.** mQTL link counts between genes inside each paralogous group. **e.** Discovered recurrent ASM loci candidate example not in HumanICR near FRG1CP for an individual with the average methylation across *self-identified Hispanic/Latino* overlay.

Examining mQTL associations across paralogous gene groups identified 305 mQTL SNP-CpG pairs on 185 paralogous genes where the associated variants are on different genes in the same paralogous group. **Fig. 5d** and **Table S12** shows the mQTL links inside each paralogous group. Across these groups, varying degrees of methylation differences occurred, indicating potential activation/deactivation patterns in a few paralogous gene groups. For example, 8 CpGs on *SCN1A* are correlated with 7 SNPs on *SCN9A* and 4 CpGs on *SCN9A* are correlated with 4 SNPs on *SCN1A*. Previous studies have shown that mutations in *SCN1A* and *SCN9A* contribute to overlapping neurological phenotypes, particularly in epilepsy and pain disorders(*77*), suggesting functional linkage between these genes. These previously unreported associations between genetic variation and methylation across SCN1A and SCN9A suggest an additional layer of regulatory complexity within paralogous gene families. mQTL density was substantially depleted among more distantly separated paralogs, particularly for pairs separated by >50 kb. Paralog pairs separated by ≤10 kb showed 0.0452 mQTLs per kb (126 mQTLs across 2.79 Mb of paralog gene sequence), compared with 0.0355 mQTLs per kb for pairs separated by 10–50 kb (148 mQTLs across 4.17 Mb) and only 0.0092 mQTLs per kb for pairs separated by 50–100 kb (31 mQTLs across 3.36 Mb), representing an approximately 80% reduction in mQTL density relative to the ≤10-kb group.

We investigated whether high F_ST_ (>0.15) variants within CAP clusters were associated with differential methylation between paralogous genes, which could suggest population specific regulation of paralogous gene copies. Overall, we identified 100 mQTL SNPs with a high F_ST_ that were associated with methylation changes in medically relevant paralogous genes. For example, mQTL links with high F_ST_ score on variants were seen in the *HLA-DQA1* and *HLA-DQB1* group (**Supplementary Fig. S29**). This could indicate population-specific regulation of these genes, potentially driven by genetic differentiation among CAP clusters. Previous studies have shown that genetic variants with high F_ST_ often contribute to phenotypic diversity and disease susceptibility(*78*), suggesting that differential regulation of paralogs through methylation may be one mechanism underlying observed differences in disease risk across populations.

For the first time we measured allele-specific methylation (ASM) in the self reported Hispanic/Latino populations(*18*) on all samples. We obtained 994 samples that were phased across at least 87.41% of genome-wide CpGs. On average, each sample contained 12,170 ASM CpGs (0.04% of all CpGs) with more than 50% of samples showing the ASM pattern. Although no parental data was available, the recurring ASMs present in ≥ 80% of *self-identified* Hispanic/Latino could indicate imprinting control regions (ICRs). To assess this claim, recurring ASM loci were intersected with known ICRs, cis-regulatory elements that mediate parent-of-origin-specific expression of imprinted genes(*18*), revealing 764 (95.6%) of recurrent ASM overlap 58.06% of HumanICR(*79*) reported loci (**Supplementary Fig. S30**). For the 24 ASM loci not overlapping HumanICR (**Table S13; Supplementary Fig. S31**), annotations from ENCODE show 17 loci overlap with cCREs, particularly PLS and pELS enriched for CTCF-binding, implicating these candidates in transcriptional regulation. Interestingly, 12 of these 24 loci, had strong support in literature as being imprinting control regions based on methylation or RNA-seq studies, including *ZDBF2*, *NDN*, *SNRPN*, *FANK1*, *ZNF595*, and *MEG8* (*80*, *81*). The remaining 12 recurring ASM were unreported and were close to *LINC01783*, *ANKRD36C, RARRES2P1* (also a CTCF-bound region according to ENCODE cCRE), *ZNG1F*, *DUX4L50*, and *LINC03025 (***Table S13)**. For example, **Fig. 5e** shows haplotype specific methylation changes closely located to the lcRNA gene *FRG1CP* where the average DNA methylation shows an allele-specific pattern.

As such our utilization of long-reads in a CAP cluster setting to study methylation represents the most comprehensive study of CpG methylation entailing novel insights across mQTLs, especially tandem repeats, and highlighting intriguing interplays of CAP cluster specific activation or deactivation of paralogous genes that were not seen before.

## Discussion

This study presents the most comprehensive genomic and epigenomic analysis of self-identified Hispanic/Latino populations in the US to date, capturing all forms of genetic variants (SNV, InDel, TR and SV) along with CpG methylation from 1,490 individuals. We reveal over 10 million novel alleles across SNV and SV alone with additional 33 million fully resolved TR alleles. By determining continental ancestry globally and further delineating local ancestry, we demonstrate that genetic diversity within self-identified Hispanic/Latino populations is both profound and structured. This has important implications for variant interpretation and disease risk assessment, and may support more inclusive genomic approaches to precision medicine. Importantly, we show that medically relevant alleles, including pathogenic and likely pathogenic variants, substantially differ in frequency across CAP clusters. The large number of SV of unknown significance highlights a lack of exploration of their impact. Moreover, our integrated assessment of SNV, InDel, SV and TR variants reveals that each variant type contributes distinctively to both genomic diversity and disease relevant loci often explained by local ancestry. These findings reiterate that self-identified race and/or ethnicity should not be used as a stand-in for genetic ancestry and demonstrate that treating “Hispanic/Latino” as a monolithic label in clinical genetics is insufficient, highlighting the need for finer resolution and ancestry aware analysis in admixed populations.

In the local ancestry inference (LAI) experiments, we tested two phasing approaches: read-based individual phasing (rLAI) and population-based phasing (pLAI). These approaches somewhat agreed on the number of ancestry switches per-sample except for genomes with a stronger African component, namely the AFR, AFR/EUR, and ADMX CAP clusters. This discrepancy exposes the complexity of phasing, especially in regions of high genetic diversity or ancestral transitions. African populations, with their shorter haplotype blocks and higher recombination rates(*82*), have greater complexity in haplotype resolution, which can result in more frequent phase errors during ancestry inference. rLAI’s agreement to the orthogonally derived haplotype-resolved long-read assembly LAI results indicates LRS may lead to a better, but more complex, phasing process, where increased switches more accurately reflect the intricate genetic landscape of admixed populations.

By integrating all types of variants (SNV, SV, TR) with base-resolution CpG methylation, we present the first comprehensive map of mQTL across all variant types(*18*). Further, this is the first long-read TR-mQTL study with detection of over twelve thousand allele-specific methylation loci from the same sequencing assay. These findings extend the significance of TR beyond their established roles in genome instability and repeat-expansion disorders, identifying repeat length as an important correlate of local DNA methylation across the population. Furthermore, we identified 24 recurrent ASM loci not reported in the HumanICR database. The improved mappability of long-reads to investigate paralogous genes revealed a novel mechanism of ancestry-linked epigenetic divergence at regulatory regions and paralogous loci, highlighting population-specific layers of gene control previously obscured by short-read approaches.

Together, our findings underscore that the power of long-read multi-omics lies not just in technological novelty, but in the unparalleled biological resolution it affords, in particular when applied to populations underrepresented in genomic research. Our results demonstrate how studying genetically diverse populations can reveal previously undercharacterized alleles, genetic–epigenetic associations, and disease-associated loci that may be missed by studies focused predominantly on European ancestry. Including diverse genetic backgrounds improves target discovery for drug development, as certain variants may only appear or exert functional consequences within specific ancestral lineages(*10*). Importantly, integration with EHR-derived phenotypes demonstrated the clinical relevance of the SV callset: SVs were included in the 80% causal set for 86.9% of prioritized associations and had the highest posterior probability in 60.0%. This is consistent with previous population-scale long-read analyses in All of Us, where SVs represented the strongest fine-mapped variant at 70.8% of associated traits in self-reported African Americans(*14*). Thus, highlighting the substantial contribution of SV to disease-associated loci that may be incompletely captured by SNV-focused analyses. Local ancestry can modulate variant effects, as seen in the *APOE4* allele, where Alzheimer’s risk is attenuated on African ancestry haplotypes compared to those from European ancestry(*83*). In precision medicine, these dynamics mean that polygenic risk scores, pharmacogenetic markers, and diagnostic thresholds can vary substantially by ancestry, and failing to account for this can lead to misdiagnosis or ineffective treatment, especially in individuals with highly admixed genetic ancestry. This study offers ancestry-specific allele frequencies, complex and large variants, and epigenetic regulatory insights, all of which can inform more inclusive and effective genomic medicine and enable biomedical discovery when coupled with clinical and phenotypic data in the All of Us resource.

Nevertheless, this study is not without limitations. Although our dataset is among the largest long-read efforts to date, larger sample sizes will be required to robustly associate rare variants with diseases and environmental factors. Additionally, while long-reads have many benefits, including simultaneous measurement of genetic and epigenetic variability with high accuracy in most variant types, the diminished performance of small InDels (1-5bp) was unfortunate. We anticipate future advances in LRS will greatly improve on this performance, including the possibility of the routine construction of complete telomere-to-telomere genomes with near perfect accuracy. Furthermore, methylation profiling in blood may not fully capture tissue-specific regulatory variation. While blood is the most often collected for research and medical studies of individuals, it has limitations to predict changes in other tissue types. Future studies integrating transcriptomic and clinical phenotypes across tissues will be crucial for deepening biological interpretation and clinical translation of methylation. Participation in All of Us is voluntary, and the geographic representation of self-identified Hispanic/Latino participants varies across U.S. states; therefore, this cohort should not be considered representative of all Hispanic/Latino individuals in the United States. Increasing the diversity of genomic datasets is only one component of improving health equity; equitable implementation and health outcomes also depend on social, structural, environmental, and healthcare factors beyond genetics.

In conclusion, our findings provide a foundational resource of admixed populations and illustrate how comprehensive, population-specific, multilayered genomic data can broaden discovery and improve the inclusiveness of precision medicine research. By revealing novel variants, regulatory mechanisms, and ancestry-specific risk factors, this work fulfills the All of Us Research Program’s mission to support genomic research that serves every community. More broadly, our findings demonstrate that integrating genetic variation, ancestry, and DNA methylation reveals layers of human genomic diversity and disease-associated variation that remain difficult to characterize using conventional sequencing approaches.

## Methods

### Sample selection

Self-identified Hispanic/Latino samples (identified in All of Us questionnaire) were chosen from release AoU v7 based on the available short read based SNV calls. Self-identified ethnicity was used for cohort selection only; genetic ancestry was inferred independently from genomic data as described below. Because self-identified ethnicity is not a proxy for genetic ancestry, this sampling strategy enriches for a socially defined group rather than a genetically discrete population; genetic ancestry was therefore inferred independently from genomic data. The sample selection for self-identified Hispanic/Latino samples was based on utmos (https://github.com/ACEnglish/utmos) a reimplementation of SVCollector(*21*) to improve speed in the process. The samples were selected based on SNV and indels on chromosome 21 to maximize diversity. As input we filtered SNV and INDEL with a minimum allele frequency of 1% using BCFtools. This output was converted to a compressed, intermediate format with utmos (--no-singelton –lowmem). Subsequently, utmos select was run with weighted allele frequencies. Here variants are weighted based on their allele frequency to emphasize the selection of the most common but diverse variants. The sample selection was done to avoid overlap with previous AoU selections for LRS.

### Paralogous gene catalog

The paralogous gene catalog was compiled using Ensembl Biomart (https://useast.ensembl.org/info/data/biomart/). Ensembl Genes 113 was selected as the main database for Humans (GRCh38.p14). In the Attributes tab, from “GENE” subcategory the “Homologues” option was selected, with “Chromosome/scaffold name”, “Gene start (bp)”, “Gene end (bp)” and “Gene name”. From the “PARALOGUES”; “Human paralogue associated gene name”, “Human paralogue chromosome/scaffold name”, “Human paralogue chromosome/scaffold start (bp)”, “Human paralogue chromosome/scaffold end (bp)” and “Paralogue %id. target Human gene identical to query gene” were selected. For the “Filters” category, chromosomes 1-22, X, Y were selected and “Paralogous Human Genes: Only”. On the Results tab, “Unique results only” option was checked and TSV output was generated. Output was processed to select best matches with score >40 and remove duplicated entries using dedup.py script.

### Nanopore sequencing

#### Baylor College of Medicine Pipeline

Genomic DNA extracted from whole blood was quantified using the Lunatic UV/Vis spectrophotometer (Unchained Labs). DNA size was determined using the Agilent Femtopulse. DNA was diluted to 60 ng/µL using nuclease free water to a volume of 50 µL. Covaris g-tubes (Covaris 520079) were used for shearing starting with 3 µg of the diluted DNA. Each sample was passed four times through the orifice of the g-tube using centrifugation at 3,800 rpm. Following bead purification, using AMPure XP beads (Beckman Coulter), the sheared DNA was size-selected on the PippinHT instrument (Sage Science) using the 6-10 kb High-Pass definition with a minimum size selection threshold of 10 kb. Loading and elution of samples from the PippinHT cassette were performed following manufacturer instructions. The eluted sample was then bead purified using AMPure XP beads (Beckman Coulter) and resuspended in 50 µL nuclease free water. The average insert size was determined by analyzing the eluate (1 µL) on the Agilent 2100 Bioanalyzer using the DNA 12000 chip for an expected average size of 15-20 kb. Modifications to this protocol were made to achieve 25-30 kb average library sizes: Starting with 7.5 µg, genomic DNA at 80 ng/ul was sheared using g-tubes using 2 passes at 3,800 rpm. Size selection was performed using the 15-20 kb High-Pass definition on the PippinHT with a threshold of 20 kb. Subsequent steps were identical for both library sizes. An aliquot (48 µL) of the purified fragments was then used as input for the ONT SQK-LSK114 library preparation kit. End repair/damage repair and adapter ligation were performed as per manufacturer’s instructions. Post ligation, libraries were purified using 1X AMPure XP beads and washed using the Long Fragment Buffer. Final libraries were eluted in 27 µL of the ONT elution buffer and quantified using the Qubit dsDNA quantification broad range assay (Thermo Fisher Scientific). ONT sequencing libraries were processed using R10.4.1 flow cells (FLO-PRO114M, ONT) on a PromethION device (PromethION 24, ONT) with default settings. Briefly, 150-200 ng of sequencing libraries, with an average insert size of 15-20 kb, were loaded onto the flow cell. Sequencing data were collected for 80 hours using MinKnow v22.10.7. On-board basecalling was performed using the super-accurate (SUP) model. If nanopore blockage occurred in the flow cell, samples were nuclease washed and reloaded with 150-200 ng of the library.

#### Johns Hopkins Pipeline

Genomic DNA extracted from whole blood was quantified using the Lunatic UV/Vis spectrophotometer (Unchained Labs). 3ug of DNA was normalized to 75ng/ul in 40uls using nuclease free water and sheared with Covaris g-tubes (520104, Covaris). Each sample was passed two times through the g-tube using centrifugation at 4000 rpm for 3 min. The sheared DNA was then size selected on the BluePippin instrument (Sage Science) using the 20kb High Pass Plus Marker U1 cassette definition with a threshold of 22kb. After size selection, samples underwent a 0.8x bead based clean up (AmpureXP Beckman Coulter) and eluted in 50uLs. Samples then underwent library prep using the NEBNext Companion Module (New England Biolabs) and SQK-LSK114 library prep kit (ONT) per manufacturer’s protocol. Post ligation, libraries were purified using 0.8x bead based clean up (AmpureXP Beckman Coulter) and washed using the Long Fragment Buffer. Final libraries were quantitated using the Qubit dsDNA Broad Range assay (Thermo Fisher Scientific). Libraries were then loaded (16-25fmol) onto R10.4.1 flowcells and sequenced on a PromethION 24 device (ONT) for 72hrs with reloading of library every 24hrs for a total of 3 loads. On-board basecalling was performed using the super-accurate (SUP) model. Later additional DNA was available from the repository and the DNA input was increased to 6ug normalized to 75ng/uL in 80uLs. Protocol modifications included a 0.8x clean up post shear to reduce the volume according to the input required for size selection on the BluePippin instrument. In addition, the threshold was adjusted to 25kb using the 20kb High Pass Plus Marker U1 cassette definition. All other steps were as described.

### PacBio sequencing

#### PacBio HiFi sequencing library preparation

Genomic DNA extracted from whole blood was quantified using the Lunatic UV/Vis spectrophotometer (Unchained Labs). DNA size was determined using the Agilent Femtopulse. Starting with 7.5 µg of DNA, shearing was performed using g-tubes (Covaris Part number 520079) by centrifuging at 4,100 rpm. Each sample was passed four times through the orifice of the g-tube. Following bead purification using Pacbio SMRTbell cleanup beads, the size of the sheared DNA was determined on the Agilent Femtopulse or the Agilent 2100 Bioanalyzer using the DNA 12000 chip. The anticipated size range was 18-22 kb. Sheared DNA was size-selected on the PippinHT instrument (Sage Science) using the 15-20 kb High-Pass definition. The size selection threshold was set at 15 kb. The loading and elution of samples from the PippinHT cassette were performed following manufacturer’s instructions. The eluted sample was bead purified using Pacbio SMRTbell Clean up Beads and resuspended in 47 µL of Pacbio Elution Buffer. The Pacbio SMRTBell Prep Kit 3.0 was used for library preparation using 46 µL of this eluate. DNA damage repair, A-tailing and adapter ligation were performed as per manufacturer’s instructions. SMRTBell Adapter Index Plate 96A was used for barcoding each library. Adapter ligated DNA was nuclease treated following the manufacturer’s guidelines and purified using 1X Pacbio SMRTBell Cleanup beads. Final libraries were eluted in 30 µL Pacbio elution buffer and quantified using the Qubit dsDNA quantification high-sensitivity assay (Thermo Fisher Scientific). Final library size was determined using the Agilent Femtopulse.

#### PacBio Sequencing on PacBio Revio Instrument

SMRTbell libraries were sequenced on the PacBio Revio instrument following the manufacturer’s protocol. Briefly, SMRTbell libraries underwent SMRTbell Binding and Polymerase Complex Formation using the PacBio DNA Polymerase (Revio Polymerase Kit). Annealing and binding conditions, including primer:template ratios, were calculated using the SMRT Link Binding Calculator. Polymerase-template complexes were loaded onto Revio SMRT Cells (25 million SMRT Cells, Pacific Biosciences) at an optimized concentration/ (170 pM with SMRT V12 and 325pM with SMRTlink V13). Sequencing runs were initiated via SMRT Link using the HiFi sequencing protocol, with a 30-hour movie time per SMRT Cell to generate high-fidelity (HiFi) reads.

Raw data underwent initial processing onboard the Revio instrument. This included real-time base calling and HiFi read generation via the Circular Consensus Sequencing (CCS) algorithm, which polishes subreads into highly accurate (>99.9% accuracy) consensus sequences. Processed HiFi data were transferred to the HGSC compute cluster for further analysis. Post-processing steps included adapter trimming and sample demultiplexing, performed using PacBio’s SMRT Analysis software (v12.0, V13). Final HiFi reads were formatted for downstream applications, such as genome mapping, structural variant detection, and variant calling.

### PacBio Assembly generation and high confidence regions

Raw PacBio hifi(HIFI) reads were extracted from bam using samtools(version:1.9) with command: samtools fasta -n -@ 12 ${bam} > ${fasta} and assembled by hifiasm v0.19.6 with command: hifiasm -o ${out} ${fasta}(*84*). This generated one primary assembly and two phased assemblies for each sample. Summary metrics for assemblies including NGx values from each assembly were obtained by running Assemblie-stats (version 1.0.1) (*85*). Phased assemblies from each sample were mapped against the GRCh38 genome using minimap2(version:2.20-r1061) with parameters: -cx asm5 --secondary=no --cs. paftools.js(version:2.17-r941) was used to call variants with parameters: -q60 -L10000 and were used for benchmarking. Aligned regions were obtained from each assembly from the minimap2’s paf output. Well covered reference regions (1 alignment per-haplotype) were collected by command BEDtools genomecov -i aln.bed -g ${bref}.fai -bga > cov.bed. Any reference region which was not well covered by at least 95% of the assemblies was marked as a low confidence region for SV discovery/genotyping. The final set of SV confident regions span 2,606,864,796bp (84.4%) of GRCh38. We next intersected the 5,027 Medical Relevant Genes(*86*) with the HQ regions and found 455 genes aren’t at least partially covered by the regions, with most (269) of the uncovered genes coming from chrX. Furthermore, an additional 120 of the Challenging Medical Relevant Genes are entirely outside the HQ regions(*86*).

### Single sample processing

All individuals were processed with Princess v2(*87*) which aligns the reads to GRCh38 reference (without alternative contigs) using preset parameters from Minimap2(*88*) for ONT and reports out a bam file. We further processed control samples from HPRC. Each sample of HPRC was downsampled randomly to achieve a similar coverage to all the ONT samples from our study.

### Small Variants

#### SNV and INDEL calling

SNV calling was performed using Clair3 v3.0.1.11(*28*). To prepare calls for merging, sample names were inserted into the VCF header (samplenames.py; see GitHub repository (https://github.com/fritzsedlazeck/Aou_hispanic) for all custom scripts), based on the sample IDs. Further, headers were modified to include absent contig definitions in some samples (headercontigs.py). For consistent merging, ambiguous nucleotide symbols within GVCF callsets were masked with “N” (filternucl.sh).

SNV calls were split on a per-chromosome basis, roughly 10% of the genome in each chunk (see GitHub repository for complete set of commands used). Resulting fragments of GVCF sample files were merged using GLNexus v1.4.1(*89*) within each chunk of the genome. The GLNexus was supported with Jemalloc v5.2.1(*90*) to optimize resource utilization. The clair3.xml is the default Clair3 GLNexus configuration file, available at Clair3 author repository (http://www.bio8.cs.hku.hk/clair3_trio/config/clair3.yml).

LD_PRELOAD=“libjemalloc.so” glnexus_cli \

-c clair3.xml \
--mem-gbytes 96 \
--threads 24 \
--list “$input_GVCF_list_to_merge” \
--dir “$output_dir”

For quality control, the resulting calls were further filtered for a minimum of 10% GT rate (missingness) and minimum of 15% GQ rate - genotype quality (filterBCF_gt.py script). Further, loci were normalized using BCFtools v1.19 (*91*) norm function with -m parameter, empty loci were discarded with BCFtools view (-i ‘INFO/AF>0’) and tags were filled using fill-tags. The filtered per-chromosome files were concatenated to a single GVCF file, containing all of the SNPs/InDels called over the tested cohort. GVCF files were converted to Plink format for further analysis, using Plink 2.0 (see GitHub for detailed commands on each step).

#### Small Variant Annotation

For annotating the SNVs, Annovar (version 2020-06-07) (*46*) was used on the multi-sample SNV VCF file. The following databases with respect to GRCh38 reference were used in the annotation.

- refGene (20211019) - all annotated transcripts in RefSeq gene
- avsnp151 (20170929) - dbSNP151
- dbnsfp47a(20240525) - predictive scores from dbNSFP(v3.0a)
- Clinvar_20240917 (20240924) - CLINVAR database with Variant Clinical Significance
- 1000g2015aug (20150824)
  - ALL.sites.2015_08, AFR.sites.2015_08, AMR.sites.2015_08, EAS.sites.2015_08, EUR.sites.2015_08, SAS.sites.2015_08
- exac03 (20151129) - ExAC 65000 exome allele frequency data
- Gnomad41_genome (20240602) - gnomAD genome collection (v4.1)

The “known” variants are identified in the annotation file if the avsnp151 column value is not “.” and the “novel” variants are with the avsnp151 column value equal to “.”. The “rare” variants in the population were defined if the AF < 0.01 or equal to “.” in the columns

ALL.sites.2015_08 (1000 genome database) and gnomad41_genome_AF (gnomAD) gnomAD database. For pathogenicity, we combined the ClinVar significance along with review status with REVEL and CADD_phred scores. The variants are termed pathogenic or likely pathogenic if the CLNSIG column value is “Pathogenic”, or “Likely_pathogenic”, or “Pathogenic/Likely_pathogenic” with review status higher than one gold star (reviewed by expert panel or practice guideline or criteria provided, multiple submitters, no conflicts) and REVEL_score >= 0.75 and CADD_phred >= 20.

### Structural Variants

Sniffles v2.0.5 was run to discover SVs per-sample (*25*). Variants (50-10kbp insertion and deletions) were then consolidated using BCFtools v1.19 (*91*) and collapsed using Truvari version 5.2 (*25*, *26*). Kanpig(*27*) v1.1 was run to genotype SVs with parameters --ploidy-bed –hapsim 0.97 --neighdist 500 --maxpaths 1000 --seqsim 0.85 --pileupmax 30 --gpenalty 0.03 --squish --ab 0.2. Results were then merged with BCFtools, preserving only vcf entries present in at least one sample after masking all alternate allele genotypes with less than 3x coverage as missing. Annotations were added using BCFtools +fill-tags, and Truvari annotations “addid, grpaf, gtcnt, lcr”. SVs which did not have both breakpoints within a single entry of the ARD regions (Methods PacBio Assembly generation and high confidence regions) had their FILTER set to ARD.

#### SV Annotation

SVAFotate(*44*) was used to intersect SVs to prior sets of discovered SVs from TopMed, gnomAD, CCDG, and 1kGP. With parameters -f 0.8 -a best. SVs with at least one intersection to any of the four reference SV databases were considered to be known. Allele frequency correlations calculated with a Pearsonr and the reference SV dataset’s reported AF from the SV with the highest reciprocal overlap to the AoU SV.

Additionally STIX (*45*) (de50db8) population frequency annotations were added using parameters -i {giggle_index} -d {stix_index} -f {vcf} -s 100 -T 5 > {out_vcf}. The indexes were built based on the 1000 Genome Project (N=1,008) and Human Pangenome Reference Consortium (N=100).

AnnotSV(*92*) (v3.4.4) was used to annotate the structural variants with the following command and parameters.

AnnotSV \
-annotationsDir “$Annot_Dir” \
-variantconvertDir “$variantconvert_Dir” \
-annotationMode “both” \
-benignAF 0.01 \
-candidateGenesFiltering 0 \
-genomeBuild “GRCh38” \
-includeCI 1 \
-metrics “us” \
-minTotalNumber 500 \
-overlap 100 \
-overwrite 1 \
-promoterSize 500 \
-rankFiltering “1-5,NA” \
-reciprocal 0 \
-snvindelPASS 0 \
-SVinputInfo 1 \
-SVminSize 50 \
-variantconvertMode “full” \
-vcf 1 \
-SVinputFile “$input_vcf” \
-outputFile “$output_file”

For predicting the functional consequence of structural variants, CADD-SV(*93*) (v1.1) was used with some preprocessing steps. The first preprocessing step was to exclude the BND and CNV types and only keep the PASS variants of INS, DEL and DUP types. Then the VCF file was converted to BED regions which was used as an input to the CADD-SV. The singularity image was used to run the CADD-SV and the following are the commands and parameters.

singularity run \

--bind ${base_dir}/CADD_input:/app/CADD-SV/input \
--bind ${base_dir}/CADD_output:/app/CADD-SV/output \
--bind ${base_dir}/CADD_workdir:/app/CADD-SV/.snakemake \
--bind ${base_dir}/CADDSV/CADD-SV/annotations:/app/CADD-SV/annotations \
--bind

${base_dir}/CADDSV/CADD-SV/annotations/genomegitar:/app/CADD-SV/annotations/genomeg itar \

--bind ${base_dir}/CADD_workdir:/app/CADD-SV/beds \
--bind ${BIND_TMPDIR}:/tmp \
--writable-tmpfs \
${base_dir}/cadd-sv.sif \
-a /app/CADD-SV/input/v0.2.2/chunks/${chunk_name}.bed.gz
/app/CADD-SV/output/v0.2.2/${chunk_name}_score.tsv.gz

Parsing of the CADD, SVAFotate, and STIX annotation was done with custom scripts present in the project’s GitHub repository (https://github.com/fritzsedlazeck/Aou_hispanic) under sv_annotation. We further investigated SVs which ACMG classification was pathogenic or likely pathogenic. For these SVs we considered them as “potentially functional impact” if:

1. The SV has at least ten allele counts
2. The SV has homozygous alternative genotype in at least one carrier
3. The SV is not present in any of the control samples from HPRC. This is filters can be found in get_GT_vcf_w_sample_names.py

The rest of the SVs were just described.

#### Transposable Element Annotation

To annotate the SV catalog for presence of transposable elements, RepeatMasker (4.1.7-p1) was used with the parameters RepeatMasker -s -e hmmer -pa 24 -species homo_sapiens <input>. The output was annotated using “One code to find them all” (OCTFA) (*94*) tool for collapsing and parsing individual TEs. Similarly, the reference genome TE annotation was parsed. INS SVs were subsetted with tandem repeat addoto catalog, to focus on non-TR elements. The OCTFTA reference annotations were intersected with reported DELs using bedtools. Further, the TE mediated DELs were selected using find_nahr.py script. INS were annotated using ins_te_covered.py. Additionally, SVA element-associated output was selected from unprocessed RepeatMasker output (grep SVA RM.out), since OCTFA does not process them. They have been processed with find_nahr.py script and annotator_nahr.py was used to add NAHR SVA events to the database. Python script searchSVAins.py was used to select results of conserved SVA regions from the RepeatMasker result, and annotator_insdel.py script was used to add SVA annotations to the database.

### Tandem Repeat Variants

The adotto TR catalog v1.1(*63*) was intersected with PacBio assembly derived variant and TR regions which contained at least one ≥5bp InDel or SV were retained in order to generate a set of TR regions over which TR calling would be performed.

We ran medaka v2.1 (https://github.com/nanoporetech/medaka) on ONT samples and TRGT v1.0.0(*66*) on PacBio samples. For ONT we used medaka tandem --threads {threads} --phasing prephased --model {basecaller_model} --ignore_read_groups /bam/{bam_file_name} /ref/{reference_name} /bed/{tr_catalog} {sex} /out with basecaller model set as ‘dna_r10.4.1_e8.2_sup@v3.5.1’ and ‘dna_r10.4.1_e8.2_400bps_sup@v4.2.0’ respectively. All resulting VCFs were converted and merged into tandem-repeat databases using TDB (v0.3).

### Variant Benchmarking

For SNPs and small InDels, vcfdist(*95*) v2.5.3 was used to compare the 421 ONT samples’ small variants (≤49bp) against the PacBio replicates’ assemblies (see above). A bed file constructed from the intersection of the high confidence regions per-sample (1x aligned coverage per-haplotype) and the 95% high confidence regions. Only PASS variants within these regions were evaluated.

For structural variants, truvari(*26*) (v5.2.1) was used to intersect the discovered SVs (50bp to 10kbp) to the assembly derived SVs using parameters --sizemax 10000 --no-ref a --pick multi --passonly --refine. Each sample’s present (non-reference homozygous, non-missing) variants were compared independently to their corresponding assembly’s present variants. Only regions which were confidently covered (1x per-haplotype) by the assemblies and by 95% of all assemblies were analyzed.

For tandem repeats, truvari was again used to compare the TR alleles per-sample to their respective assemblies. Only TR regions which were analyzed by the TR caller (See Methods TR) which were fully spanned by each sample’s assembly high confidence region and by 95% of all assemblies as described in (see Methods: high confidence regions) were analyzed. The truvari commands used were a two step process which replicates the TR benchmarking process recommended in English et al. (*63*) of truvari bench --sizemin 5 --pick ac --no-ref a and truvari refine -w -u -c O.

### Population Analysis

#### Principal Component Analysis

Plink2 was run to perform the principal component analysis on small variants (SMs) and SVs with parameters --geno 0.05 --hwe 0.001 --maf 0.05 --pca 10. SM and SV variants were pre-filtered for those outside of linkage disequilibrium which was again calculated with plink2 and parameters --indep-pairwise 100 10 0.1. Only SVs with a PASS filter as well as those which passed the batch tests described below. Procrustes analysis of SV and SM PCAs for distance comparison and plotting was performed with scipy v1.10.1(*96*). Batch effect SVs (**Supplementary Section 4.7**) were determined by using a Gaussian Mixture Model (scikit-learn v1.6.1 (*96*)) with k=2 over the third principal component for all PASS SVs, thus clustering the samples into two batches corresponding to ONT basecaller version. Variants with a Hudson F_ST_ (*97*) ≥ 0.02 between the two batches were excluded for subsequent PCAs.

#### Admixture Analysis

The gnomAD reference panel(*98*) comprising HGDP and 1kGP samples with known super populations were merged with the AoU small variants (filtered for AF ≥ 0.05, missingness ≤ 0.05). ADMIXTURE(*31*) v1.3.0 was then run with K=3, for both unsupervised and supervised modes. In supervised mode, an additional metadata file was provided with information of super-populations for HGDP and 1kGP samples. Afterwards, the proportion columns for unsupervised results were labeled based on which were maximized for each group of the known super-populations of Amerindegenous (AMR), African (AFR), and European (EUR).

AoU continental ancestry labels were assigned using a two-majority strategy capturing at least 75% of the super-population contribution. For example, if AMR accounted for 80% of a sample’s ancestry, the sample was labeled AMR, as 80% exceeds the 75% threshold. If EUR contributed 60% and AMR 30%, the sample was labeled AMR,EUR. Samples for which no two super-populations collectively reached ≥75% were labeled ADMX.

To replicate the super population contributions and labeling results, we also analyzed the AoU variants over previously published ancestry informative markers (AIMs) (*32*, *33*). The two sets of AIMs were consolidated and deduplicated after being pulled with GRCh38 coordinates using ensembl (*99*). Only AIMs with major/minor nucleotides as reported from the original studies that matched the nucleotides after liftover were kept, resulting in a total of 540 markers. Each AIM was then pulled from 1kGP and HGDP vcfs from https://ftp.1000genomes.ebi.ac.uk/vol1/ftp/data_collections/1000G_2504_high_coverage/working/20220422_3202_phased_SNV_INDEL_SV/1kGP_high_coverage_Illumina.chr*.filtered.SNV_INDEL_SV_phased_panel.vcf.gz and ftp://ngs.sanger.ac.uk/production/hgdp/hgdp_wgs.20190516//hgdp_wgs.20190516.full.chr*.vcf.gz. HGDP AIMs were then imputed/phased using Beagle (*100*) v5.5 and the 1kGP markers as the reference panel. The same Beagle procedure was run on the AIM sites’ genotypes for the AoU samples using the merged HGDP+1kGP as the reference panel. The genotypes from AoU samples and HGDP+1kGP samples with AMR, EUR, or AFR known super population were then pulled and formatted to be run with ADMIXTURE v1.3 and K=3.

#### Local Ancestry Inference

To define specific ancestry regions we first phased the small variants with two strategies. The analysed samples were individually phased using Whatshap(*101*) (2.3) and merged together using BCFtools, per chromosome. The merged calls were annotated with BCFtools to add genotype quality, depth, allele frequency and genotype likelihood tags. Further, BCFtools was used to ensure correct reference genome sequences within calls and decompose multiallelic records. The non-phased REF/ALT variant genotypes (non-informative for phase-based ancestry inference) were converted to reference (0/0 genotype, custom_gtfix.py script) and saved in compressed vcf format. Finally, the per-chromosome files were merged using BCFtools merge and missing genotypes were set to 0/0 (required for RFmix to work correctly), recalculating allele frequencies after all the edits, both using BCFtools. Variants under 1% population allele frequency were discarded for quality-control.

Alternate analysis strategy was based on population phasing using ShapeIT5 v5.1.1 and high-coverage 1kGP Illumina SNV reference panel(*102*). The GLNexus-merged individual SNV/InDel calls (see SNV and INDEL calling methods section) were used as input for the phase_common algorithm of ShapeIT5. The analysed variants were limited to those occurring at >=0.1% population VAF, according to the software manual (*35*).

To examine ancestry regions in detail across phased small variants, we employed local ancestry inference, based on 1kGP/HGDP(*23*, *30*) reference panel, using RFmix v2 for both population and individual-based phasing approaches. Finally, perfect flips between haplotypes were corrected using a modification to the ‘unkinking’ procedure described by Atkinson et.al.(*36*). This procedure simply finds ancestry blocks that are directly adjacent if flipped between haplotypes would increase the per-ancestry continuity.

For all custom code and commands used, refer to GitHub (https://github.com/fritzsedlazeck/Aou_hispanic).

### Transient SV analysis

Using the CAP cluster classification, we annotated SVs that appear in a single individual of a given cluster (i.e AMR) but that are more widely observed (over 10% of individuals) in at least one different cluster (i.e AFR). We denoted these SVs as singletons. We performed filtering over the singleton to remove: 1) SVs that only appear as singletons, meaning per group that have a single observation and 2) removed group cross-talk as follows:

- for the AMR group we remove all SVs which appear in the AMR/EUR group
- for the AMR/EUR we remove all SVs which appear in either the AMR or EUR group
- for the EUR we remove all SVs which appear simultaneously in the AMR/EUR and AFR/EUR group
- for the AFR/EUR we remove all SVs which appear in either the AFR or EUR group
- for the AFR we remove all SVs which appear in the AFR/EUR

The ADMX and AFR/AMR groups were ignored due to their low number of samples present, furthermore, the HPRC samples with EAS and SAS were not counted. Finally, we removed one SV that was a singleton, but in one of the control samples (HPRC). The sex chromosomes were ignored from the final SVs dataset as their local ancestry cannot be assessed.

The final selection of SVs was denoted as transient SVs and was annotated with CADD, STIX, Tandem Repeats (TR) and Transposable Elements (TE), and Medical Relevant Genes.

Next, we analyzed TE-mediated SV which AF < 1% (labeled as sparse SVs), as a second source of transient SVs. We only used SVs that were present in at least two CAP clusters. Next, we performed an ancestry summarization of each SV by counting the proportion of alleles from the three main continental ancestries: Amerindegenous (AMR), African (AFR), and European (EUR) over all carrier individuals. For example, if an SV is present in two individuals whose ancestry is AMR/EUR and AMR/AFR, the ancestry summarization is as follows: AFR: 25%, AMR: 50% and EUR 25%. This summarization was done both based on the CAP cluster and Local Ancestry Inference (LAI). Previous results from the singleton analysis showed that the LAI showed a much fine grained ancestry, thus we favored the LAI summarization for the rest of the analysis. In the ancestry summarization, each ancestry component represents a “pull” towards the nodes of an equilateral triangle. This way, we were able to project the summarization of the three continental ancestries into 2D-space. We measured the Euclidean distance of each SV that has at least five carriers to the Null, which is represented by the average ancestry over all individuals based on CAP clusters. We ordered the SVs based on this distance and manually curated the top 100 SVs using samplot plots. The curated SVs are considered as transient SVs. We annotated the first 10 transient SVs with CADD and Medical Relevant Genes.

### Phenotypic and Pathogenetic TR analysis

The catalog of 187 TRs with previously reported phenotypic and pathogenic impacts were obtained from https://github.com/ACEnglish/adotto/blob/main/pathogenic/Patho.tsv, where we picked the intersection of it and our adotto catalog. There are several entries in the phenotypic catalog that cover multiple entries in the adotto catalog, where we later ignored those. We further intersected this phenotypic catalog with strchive (strchive.org) to get 61 pathogenetic TRs. We then performed the One-way ANOVA (Analysis of Variance) analysis between the TR length delta (TR length - reference TR length) and the CAP cluster. The ANOVA was performed with stats models package from python using formula-based Ordinary Least Squares (OLS) regression. We visualized the pathogenetic TR example with motifscope(*103*) to manually check and visualize all the pathogenetic tandem repeats.

### Phenome-wide SV analysis

To investigate the clinical relevance of SVs, we evaluated SV-phenotype associations in 1,490 ONT-sequenced AoU participants. We excluded seven related individuals with kinship coefficients >0.177 to minimize confounding due to relatedness, and restricted the PheWAS to 1,095 participants with standardized disease phenotypes available in AoU CDR v8. Association testing was limited to SVs carried by at least 20 participants. To reduce potential genotyping ambiguity at complex multiallelic loci, we excluded loci containing multiple alleles with ≥70% similarity in both sequence and length. Clinical conditions were derived from AoU CDR v8, and only phenotypes with at least 50 cases and 50 controls were retained. We retained phenotypes with at least 50 cases and 50 controls and excluded overly broad conditions or those unlikely to have a genetic basis.

Association testing was performed using Firth penalized logistic regression implemented in PLINK2 (*104*). For analyses including all participants, models were adjusted for age, sex, and the first 10 genetic principal components (PCs). For sex-specific phenotypes, analyses were performed separately within the corresponding sex and adjusted for age and the first 10 genetic PCs. Association results were retained only when PLINK2 reported successful model fitting (ERRCODE = “.”).

Given the modest size of the long-read cohort, this analysis was designed for discovery rather than definitive association testing. We therefore used p < 1 x 10^-5^ as a prioritization threshold to identify candidate SV-disease associations for locus-level fine-mapping. To further assess whether associated SVs represented leading signals at their loci, SNVs within 100 kb of each SV were tested for association with the corresponding disease phenotype using Firth logistic regression. Fine-mapping was then performed with CAVIAR (*105*) by jointly modeling each SV and the top 1,000 nearby SNVs with the smallest association p values to estimate causal posterior probabilities. We assessed SV inclusion in the 80% causal set and whether the SV had the highest posterior probability at the locus. For functional annotation, associated SVs were intersected with GENCODE v44 gene annotations (*106*) to identify overlapping genes.

### DNA Methylation

DNA methylation signals from ONT sequenced samples are obtained with Modkit v0.3.1 ‘modkit pileup --ref {reference_genome} --preset traditional --filter-threshold 0.9 {bam_file} {methylbed_file} ’. DNA methylation is piled up both on two haplotypes (by adding ‘--partition-tag HP’) and on the entire sample that does not distinguish the haplotypes. PCA was performed on all the methylation signals by filtering the coverage on the CpG location at least 5x and the CpG location is dropped if there are more than 10% of the samples that do not cover that location.

The blood autosomal DNA methylation catalog is generated with several statistics: 1. The average DNA methylation beta value across the 1,490 samples. We have profiled the DNA methylation on the CpGs that covered more than 95% of the samples, which is the same for downstream mQTL study and plink filter. With this threshold, we are able to capture 26,274,946 CpG locations on hg38. 2. Various statistical profiles on each CpG location across the samples. Besides the mean methylation value, we also listed the standard deviation (STD = sqrt((1/n) * Σ(xᵢ - x̄)²)), Interquartile Range (IQR = Q3 - Q1), Median Absolute Deviation (MAD= median(|xᵢ - median(x)|)) and Coefficient of Variation (CV= (STD / Mean) × 100%).

Based on this catalog, we are able to detect the highly variable regions across hg38 across the self-identified Hispanic populations. We chose the IQRs as the indicator of the variability of the DNA methylation level. We further binned the genome with 200 bps bins where we select the bins following criteria: 1. The IQR of low methylation (methylation level < 20) is less than 5, middle methylation (methylation level between 20 and 80)’s IQR is less than 15 and high methylation IQR (methylation level > 80)’s IQR is less than 10. Bins with more than 2 CpGs and more than 10% of those CpGs are “variable” are selected as variable regions. The neighboring bins are merged. We then overlapped the bins to encode cCRE database for meaningful region screening. We report the 200 bps bins where it overlaps with cCRE database for more than a half of the annotated regions’ length.

We later used this catalog to research the paralogous genes’ methylation status. We first filtered paralogous genes by their sequence similarity >60% and grouped those genes into groups with the python networkx(*107*) package. Within each paralogous group, we averaged each CpG sites’ DNA methylation level on each gene and compared it within the group.

DNA methylation-age association was performed with package pyaging(*108*) v0.1.13. We first obtained the Illumina EPIC V2’s contig coordinate on GRCh38 (https://zwdzwd.github.io/InfiniumAnnotation), and then obtained the correlation between the Horvath clock and the real reported age. We further ran wgbstools(*109*) and UXM(*71*) for cell type deconvolution. We first obtained the DNA methylation atlas (Atlas.U25.l4.hg38.full.tsv) from UXM GitHub repo (https://github.com/nloyfer/UXM_deconv) and used wgbstools to extract read methylation information as .pat file with command “wgbstools bam2pat -np -L {atlas_bed} -@ {threads} --genome hg38 --force --no_beta --out_dir {output_path} {bam_file}”. Then we followed up a deconvolution step with the uxm with command as “uxm deconv --atlas {atlas_file} –output {output_path} --threads {threads} {input_path}/*.pat.gz”. The output deconvolution result contains the estimated cell proportions of 40 cell types and we later picked 4 that are known to belong to whole blood WGS and show significance in all the samples. We then plotted the correlation between the age and those cell types.

We further performed three mQTL analyses where they act in cis (within +-100k bps). The mQTLs are predicting cis-acting CpGs against SNPs, SVs and TRs. ONT claims the DNA methylation calls in the read levels could reach to 98.8%(*110*). Therefore, our pre-filtering of DNA methylation signals is solely based on the sequencing coverage. Based on Ziller et al.’s(*111*) recommendation of CpG coverage to distinguish biological replicates, we used 5X for DNA methylation mQTL. As Du et al.(*112*, *113*) suggested, the M value (i.e., log2(beta/(1-beta)) is more practical to perform statistical analysis such as QTL studies. We have further transformed beta values into M values. For each chromosome, we filtered out the CpG locations with no methylation call over 5 samples, and samples with more than 20 missing CpGs after the CpG filtering. The mQTL is performed with TensorQTL (*114*), which is a GPU accelerated version of fastqtl(*115*). We first ran the entire all-versus-all CpG and variant association call with command “cis.map_nominal(genotype_df, variant_df, phenotype_df, phenotype_pos_df, prefix, covariates_df, output_dir=’.’)”. Based on this we further filtered out the variants that have no methylation signal related and CpG locations that have no variant related. Then we finally performed cis-mQTL with command “cis.map_cis(genotype_df, variant_df, phenotype_df, phenotype_pos_df, covariates_df)” to get the final mQTL result for each chromosome. We then performed FDR correction with “tensorqtl.calculate_qvalues(cis_df, qvalue_lambda=0.85)” and then obtained the q-value (FDR corrected p-value) of the variant-CpG pairs. The FDR procedure was described by Storey and Tibshirani (*116*).

The TR mQTL was performed with a simple linear regression between the length of the TRs and CpG values in cis window. Due to the compute memory and time constraint, we are only able to keep the cis window to 10kbp. We used the python statsmodels package (https://www.statsmodels.org/) for the linear regression with the exact same Sex, basecaller model and TR-PCA computed like SNP and SV as covariate input. We selected the top 5% of the most variable TRs based on their length standard deviations for the TR-mQTL analysis. We further filtered the results based on the p-value less than 0.05, minimum absolute effect size (abs(coef) > 0.1), r2 value (reflect the variance of CpG methylation explained by the TRs and covariants) > 0.1 and minimum sample number > 500.

The allele-specific DNA methylation was called by modkit based on pile up files per haplotype from phased BAM file with command “modkit dmr pair -a {hap1_bed} -b {hap2_bed}

-o {asm_bed} --segment {asm_segment_bed} --ref {reference_genome} --base C”. To systematically assess ASM loci across the genome, we examined the ASM calls from 1,490 samples and selected loci that exhibited MAP-based significance (p < 0.05) in more than 50% of individuals phased in the CpG location. Phaseblock coverage is derived from the genome phasing result by “whatshap stats –gtf {phased_vcf}”.

The ASM calling also provides automatic segmentation of differentially methylated regions (DMRs) between haplotypes. We overlapped our DMRs with previously published humanICR to cross check. The tandem repeat catalog, CpG Island and gene density are all downloaded from UCSC catalog. Novel recurring ASM loci candidates are calculated based on the modkit dmr segment result. We examined the regions with more than 80% phased samples overlapped that show as DMR regions. Novel recurrent DNA methylation loci are annotated manually with region intersection between cCRE database and gene catalogs. We manually annotated all the loci with literature search for imprinting implications and merged the close located ones.

## Acknowledgments

We want to thank Oxford Nanopore for giving us access to an early version of their Tandem Repeat calling method based on Medaka v2.1, especially Moustafa Shokrof. We would also like to thank Egor Dolzhenko, Zev Kronenberg and Daniel Agustinho for helpful discussions.

## Funding

This manuscript was supported by US National Institutes of Health (NIH) grants to Baylor/JHU (OT2 OD002751 and OD038122) and the University of Washington (OD038111). E.E.E. is an investigator of the Howard Hughes Medical Institute.

We gratefully acknowledge *All of Us* participants for their contributions, without whom this research would not have been possible. We also thank the National Institutes of Health’s All of Us Research Program for making available the participant data and samples examined in this study.

## Author Contributions

DM, HM, YH, KD, HS, SB, AS, MM, BC, BM and LM conducted sequencing and data submission. YZ, AP, JP, and ZK conducted primary analyses and managed data organization. AE, YF, LFP, MI, SB, XZ, SW, QL contributed to data analysis and interpretation. EA, GT, JH contributed to the ancestry analysis. GM, FS, RG, EV, WT, DM, KD and MCS provided advising and manuscript input. All the authors reviewed and edited the manuscript.

## Data Availability

All data sets are hosted at All of Us workbench at workbench ID: aou-rw-2e2b09d3

## Code Availability

All scripts are available at: https://github.com/fritzsedlazeck/Aou_hispanic

